# Temporal and spatial composition of the tumor microenvironment predicts response to immune checkpoint inhibition

**DOI:** 10.1101/2025.01.26.634557

**Authors:** Noah F. Greenwald, Iris Nederlof, Cameron Sowers, Daisy Yi Ding, Seongyeol Park, Alex Kong, Kathleen E. Houlahan, Sricharan Reddy Varra, Manon de Graaf, Veerle Geurts, Candace C. Liu, Jolene S. Ranek, Leonie Voorwerk, Michiel de Maaker, Adam Kagel, Erin McCaffrey, Aziz Khan, Christine Yiwen Yeh, Christine Camacho Fullaway, Zumana Khair, Yunhao Bai, Hadeesha Piyadasa, Tyler Risom, Alea Delmastro, Felix J. Hartmann, Lise Mangiante, Cristina Sotomayor-Vivas, Ton N. Schumacher, Zhicheng Ma, Marc Bosse, Marc J. van de Vijver, Robert Tibshirani, Hugo M. Horlings, Christina Curtis, Marleen Kok, Michael Angelo

## Abstract

Immune checkpoint inhibition (ICI) has fundamentally changed cancer treatment. However, only a minority of patients with metastatic triple negative breast cancer (TNBC) benefit from ICI, and the determinants of response remain largely unknown. To better understand the factors influencing patient outcome, we assembled a longitudinal cohort with tissue from multiple timepoints, including primary tumor, pre-treatment metastatic tumor, and on-treatment metastatic tumor from 117 patients treated with ICI (nivolumab) in the phase II TONIC trial. We used highly multiplexed imaging to quantify the subcellular localization of 37 proteins in each tumor. To extract meaningful information from the imaging data, we developed SpaceCat, a computational pipeline that quantifies features from imaging data such as cell density, cell diversity, spatial structure, and functional marker expression. We applied SpaceCat to 678 images from 294 tumors, generating more than 800 distinct features per tumor. Spatial features were more predictive of patient outcome, including features like the degree of mixing between cancer and immune cells, the diversity of the neighboring immune cells surrounding cancer cells, and the degree of T cell infiltration at the tumor border. Non-spatial features, including the ratio between T cell subsets and cancer cells and PD-L1 levels on myeloid cells, were also associated with patient outcome. Surprisingly, we did not identify robust predictors of response in the primary tumors. In contrast, the metastatic tumors had numerous features which predicted response. Some of these features, such as the cellular diversity at the tumor border, were shared across timepoints, but many of the features, such as T cell infiltration at the tumor border, were predictive of response at only a single timepoint. We trained multivariate models on all of the features in the dataset, finding that we could accurately predict patient outcome from the pre-treatment metastatic tumors, with improved performance using the on-treatment tumors. We validated our findings in matched bulk RNA-seq data, finding the most informative features from the on-treatment samples. Our study highlights the importance of profiling sequential tumor biopsies to understand the evolution of the tumor microenvironment, elucidating the temporal and spatial dynamics underlying patient responses and underscoring the need for further research on the prognostic role of metastatic tissue and its utility in stratifying patients for ICI.

## Introduction

Immune checkpoint inhibition (ICI) is a form of cancer immunotherapy that has initiated a paradigm shift in treatment across a range of solid cancer types^1–9^. Although ICI has demonstrated the potential for durable remission in patients who previously had few options, response rates to monotherapy have been low in patients with metastatic breast cancer^10,11^ relative to diseases like melanoma or lung cancer. Consequently, recent clinical trials have combined immunotherapy with standard chemotherapy for patients with metastatic breast cancer. The IMpasssion130-trial^12^ was the first phase III trial evaluating the efficacy of chemo-immunotherapy for metastatic triple negative breast cancer (TNBC). The KEYNOTE-355 trial found additional evidence for the efficacy of a combined chemo-immunotherapy schedule, making chemo-immunotherapy the current standard of care for patients with metastatic PD-L1-positive TNBC^13^.

Numerous studies have attempted to identify clinical, pathologic, genomic, transcriptomic, and proteomic predictors of ICI response in breast cancer^14–22^. However, it has remained challenging to identify robust biomarkers that reliably separate responders from non-responders. This is thought to be due in part to the multifaceted effects of immunotherapies. Unlike tumor-targeted therapies that are directed towards cancer cell-specific proteins, immunotherapies aim to activate the patient’s immune system. Owing to the number of distinct cell types and cell states that need to be measured to evaluate a complex immune response, assays measuring a few parameters at once have been insufficient to decode the underpinnings of immunotherapy response. In addition, the roles of these diverse cell types are often influenced by their location and interactions within the tumor microenvironment (TME)^23^. As such, assays that lack spatial information cannot fully resolve this complexity.

Cancer immunoediting occurs during the natural progression of disease, but evidence from studies using patient materials is limited. Characterizing how the TME differs in primary versus metastatic disease and how the TME correlates with patient outcome from ICI requires the collection and analysis of matched, longitudinal patient samples. To date, serial tumor sampling is rare, hindering our ability to understand the relationship between tissue dynamics, disease progression, and patient outcomes.

Here we provide a spatiotemporal dataset of TNBC, featuring matched primary tumors and longitudinal biopsies of metastatic lesions collected before and during ICI treatment (specifically anti-PD1) in the context of a prospective clinical trial. We generated multiplexed imaging data of pathology sections for each patient at different timepoints and combined this with previously generated genomics and transcriptomics^24^ data to enable multi-modal characterization of the TME. In-depth analysis of these data provides insights into how spatial proteomic, transcriptomic, and genomic characteristics evolve through disease progression and immunotherapy in TNBC and their association with patient response. Our analysis identified numerous imaging and transcriptomic features associated with response, including T cell infiltration at the cancer border and the diversity of cellular neighborhoods, as well as specific cell subtypes such as PD-L1+ myeloid cells. In contrast, we did not identify robust correlates of response from the whole-exome sequencing data generated from pre-treatment biopsies. Looking across timepoints, we found that increases in cellular diversity were consistently associated with better outcome. In contrast, other features like CD8 T cell density and PD-L1 expression levels were not universally predictive, and their association with outcome was timepoint-dependent. We constructed multivariate models to predict patient outcome, finding that on-treatment samples provided the most predictive information, whereas primary tumors exhibited almost no predictive power for ICI response. These findings underscore the importance of longitudinal sampling and multimodal characterization in defining the trajectories of patient outcome, shedding light on the determinants of ICI response and the design of future clinical trials.

## Results

### Multimodal characterization of serially sampled metastatic TNBC

We obtained longitudinal histological samples from 117 patients with metastatic TNBC who received nivolumab (anti-PD-1) in the TONIC trial (NCT02499367)^24,25^. All 117 patients were included in TONIC-I stage I (n=70, 59.8%) or stage II (n=47, 40.2%). The interval between primary diagnosis and inclusion in the TONIC trial averaged 43.8 months (range 1-284 months), while the disease-free interval from primary disease to first relapse averaged 30.5 months (range 0-237 months). Detailed clinical characteristics were collected for all patients (**Supplementary Table 1)**. To understand the evolution of the tumor microenvironment (TME), we collected tumor samples from four timepoints: 1) the primary tumor (via retrospective collection), 2) the metastatic baseline biopsy taken at enrollment in the TONIC trial, 3) the metastatic pre-nivolumab (pre-nivo) sample which immediately followed induction treatment (before ICI), and 4) the metastatic on-nivolumab (on-nivo) sample taken after three cycles of nivolumab given every two weeks (**Fig. 1a**). Samples were screened by a pathologist to identify representative regions containing tumor, and tissue microarrays were constructed with cores from each sample (**Extended Data Fig. 1a**). We designed an antibody panel to capture the diversity of cell types found in the TME, their functional states, and the core components of the extracellular matrix and tissue architecture (**Extended Data Fig. 2a, Supplementary Table 2**). To overcome the multiplexing limitations of conventional imaging techniques, we used the multiplexed ion beam imaging (MIBI) platform^26^ and a 37-plex antibody panel for quantitative subcellular imaging. By tagging each antibody with a unique elemental isotope, and using spatially-resolved mass spectrometry to read out the antibody signal, MIBI enables the acquisition of highly multiplexed imaging data without the need for cyclic staining or imaging (Methods).

**Figure 1.**
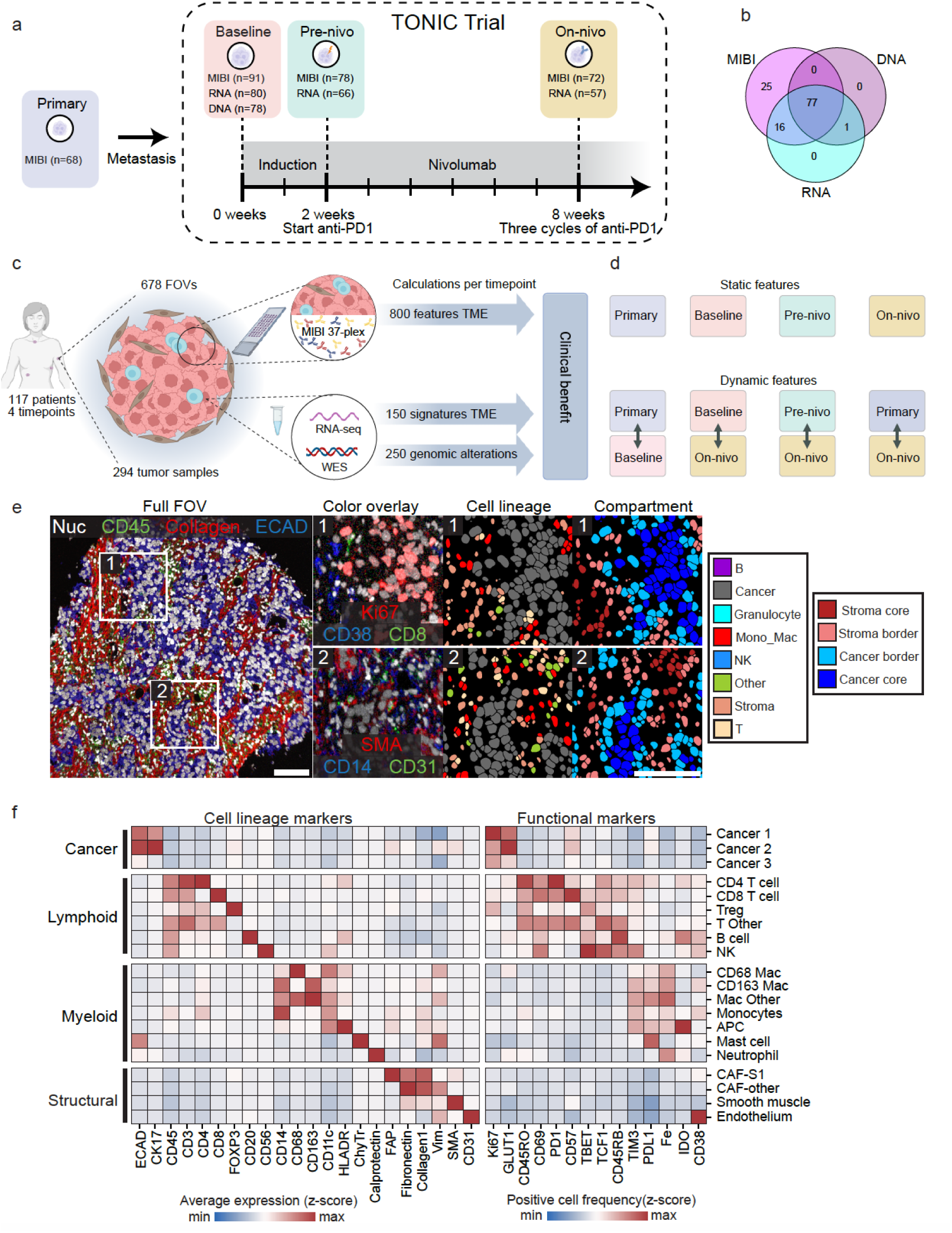
Study design, workflow and feature extraction. **a**) Schematic overview of the collection of tumor tissue for MIBI analysis from patients accrued in the TONIC trial (NCT02499367). Tissue was collected from the primary tumor, as well as metastatic samples at baseline, after induction (pre-nivo), and after three cycles of nivolumab (on-nivo). **b**) Venn diagram showing the number of patients who have overlapping data from the three modalities (DNA, RNA and MIBI) included in the study. **c**) Schematic overview of the data generation and feature extraction workflow for DNA, RNA and MIBI data. **d**) Schematic representation of static and dynamic feature data. All features were calculated and compared to response on four separate, static timepoints. In addition, the dynamic change between pairs of timepoints was also calculated and compared against patient response. **e**) Full FOV: MIBI field of view (FOV) color overlay of a representative tumor biopsy. Inset 1 and 2 show blowups at increased magnification from the original image. Color overlay: Additional color overlays from the same FOV. Cell lineage: Cell mask with cells colored by cell classification. Compartment: Cell mask with cells colored by tumor compartment. Scale bars: 100um. **f**) Heatmap showing the identified cell clusters (y-axis) with the average expression of the markers in the MIBI panel (x-axis) split by marker type. Cell lineage markers were used to generate the clusters, whereas functional markers were not used for clustering.

We generated multiplexed images from 678 distinct TMA cores (**Extended Data Fig. 1b-c**), encompassing 64 primary tumors, 91 baseline biopsies, 78 pre-nivo biopsies, and 72 on-nivo biopsies across 117 unique patients (**Fig. 1a, Extended Data Fig. 1d, Supplementary Table 3**). Raw MIBI images were background subtracted, denoised, and normalized prior to analysis (**Extended Data Fig. 3, 4**, Methods).

To complement the spatially-resolved proteomic information, we analyzed bulk whole-exome sequencing (WES) from the same metastatic baseline samples and bulk whole-transcriptome RNA sequencing (RNA-seq) from the same metastatic baseline, pre-nivo and on-nivo samples (**Fig. 1b**), using a harmonized, state-of-the-art bioinformatics pipeline (Methods). In total, we aggregated WES data from 78 patients, RNA-seq data from 203 samples across 92 patients, along with the previously described MIBI data from 305 samples across 117 patients, for a total of 117 patients profiled with any of the three modalities (**Fig. 1b-c, Extended Data Fig. 1e-g**).

Following generation and processing of the image data, we used our previously developed tools to identify and classify the cells in each image (**Fig. 1e**). Namely, for cell identification we employed Mesmer^27^, a pre-trained deep learning algorithm for automated, accurate cell segmentation (**Extended Data Fig. 5a-c)**. We identified an average of 2905 cells (range 11-11687) in each of the distinct cores (**Extended Data Fig. 5d**). For cell classification, we used Pixie^28^, an algorithm created to cluster cells in spatial datasets. This enabled us to identify 22 intermediate cell clusters (**Figure 1f**) which were binned into eight broad cell lineages summarizing the major components of the TME (**Extended Data Fig. 6a-d**).

With the location and subtype of each cell identified, we quantified the abundance of the cell clusters in the baseline pre-treatment metastatic tumors from our patient cohort. Cancer cells were the most abundant cell type, constituting 52.4% of segmented cells, divided into three groups: Cancer 1 (61.7% of total cancer cells), Cancer 2 (13.4%), and Cancer 3 (24.9%). Cancer 1 and Cancer 2 cells predominantly expressed epithelial markers E-Cadherin (ECAD) and Cytokeratin 17 (CK17), with increased proliferation (Ki67) and glucose uptake (GLUT1). Cancer 2 cells showed the highest expression of □SMA, a marker associated with cancer cell invasion and metastatic potential^29^. Cancer 3 cells were characterized by dim expression of ECAD and CK17 (**Fig. 1f, Extended Data Fig. 6g**). Immune cells were the next most abundant cell type (25.6% of all cells, **Fig. 1f, Extended Data Fig. 6h**). CD4^+^ T (7.8% of immune cells) and CD8^+^ T cells (11.1% of immune cells) were two of the most abundant immune cell populations and often expressed the immune activation marker PD-1. Tregs (7.5% of immune cells) were the most proliferative immune cell population, with 16% expressing Ki67^30^. Antigen presenting cells (APCs) exhibited the highest expression of IDO1, known for inducing long-term immunological tolerance by limiting effector T cell activity and promoting Treg induction^31,32^. We identified four distinct macrophage/monocyte populations, many of which expressed PD-L1 and TIM3. Natural killer (NK) cells, though rare (0.5% of immune cells), often expressed T-BET and TIM3, suggesting their maturation and immunosuppressive status ^33–35^.

Structural cells (21.5% of total) included fibroblasts (74% of structural), endothelial cells (16%), and smooth muscle (9%, **Extended Data Fig. 6i**). Among fibroblasts, we identified cancer-associated fibroblast (CAF) phenotypes similar to CAF-S1 (40%), previously linked to immune suppression and Treg interaction in TNBC^36^, as well as a CAF-Other population (35%) which did not express FAP or SMA. We had two clusters for uncategorized cells: immune cells that were positive for CD45 but negative for other markers (Immune Other, 3% of total), and cells that were negative for all markers in the panel (Other, 0.5% of total). Overall, the observed abundances of major cell populations align with previous spatial analyses of TNBC^37,38^.

We next examined changes in cell population prevalence in metastatic lesions from baseline to on-nivo. Baseline (n=91) and on-nivo (n=72) metastatic tumors had similar cell proportions (**Extended Data Fig. 6e**). This conservation of cell proportions across timepoints included distinct cancer cell subtypes, as well as structural, myeloid, and lymphoid populations. When using a more detailed cell clustering with 22 discrete clusters rather than eight broad cell lineages, baseline and on-nivo timepoints remained similar (**Extended Data Fig. 6f**). Here, we identified and classified the major and minor cell types across all images in our cohort. The modest differences across time when looking only at cellular abundance prompted us to develop metrics to capture the spatial dynamics in the tumor microenvironment.

### Comprehensive quantification of the tumor microenvironment with SpaceCat

Highly multiplexed image data is a rich source of information for defining the spatial relationships between distinct cell types. However, quantifying this information in an accurate and scalable manner is a significant challenge. To address this gap, we developed SpaceCat, an open-source computational pipeline which generates a spatial catalog of informative features from multiplexed image data. SpaceCat can be applied to any processed multiplexed imaging dataset to summarize the abundance, location, and phenotype of the cellular and acellular features of each image. We used SpaceCat to generate 872 features that quantify cell density, spatial relationships between cell types, extracellular matrix composition, immune infiltrate organization, functional marker expression, and cellular diversity (**Fig. 2a, Extended Data Fig. 7, Supplementary Table 4**, Methods). These features include cell density statistics

**Figure 2.**
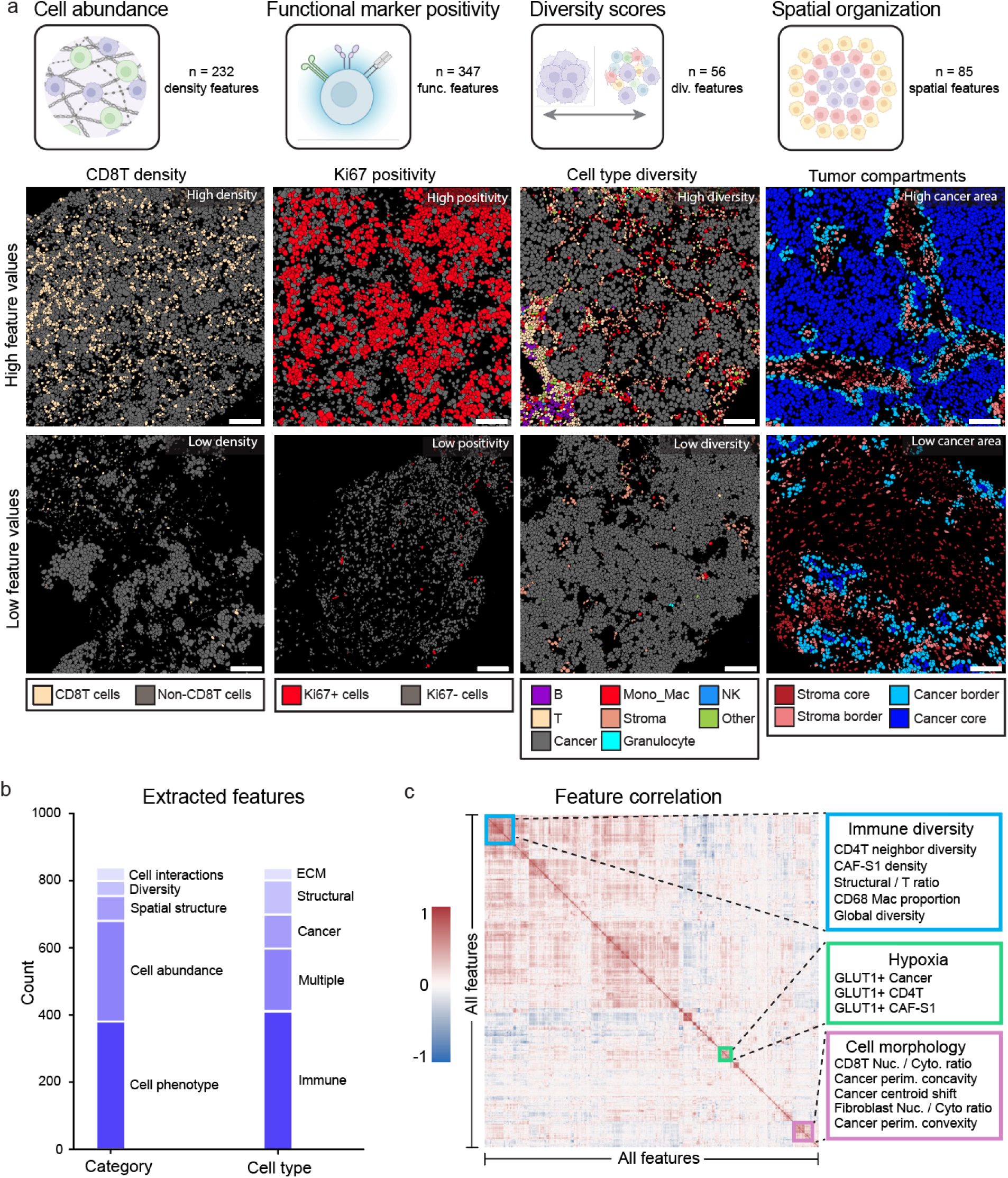
SpaceCat feature extraction pipeline. **a**) Four categories of features (cell abundance, functional marker positivity, diversity scores, and spatial organization) calculated by SpaceCat, with an example feature belonging to each of those categories. Each column shows one example image with a high value of the feature and one example image with a low value of the feature. Scale bars: 100um. **b**) Distribution of extracted features according to the category of the feature (left) and cell type associated with that feature (right). The category cell phenotype (n=453) includes features based on functional markers (n=357) and morphology (n=96). The category structural organization (n=85) includes features base on extracellular matrix (n=78) and tumor compartments (n=7). The category interactions (n=30) include features based on linear distances (n=24) and the mixing of cells (n=6). The category diversity (n=58) includes features for cellular diversity (n=38) and regional diversity (n=20). The category cell density (n=232) includes features for density (n=90), proportional density (n=49) and density ratios of cells (n=93). **c**) Clustered pairwise correlation of features across all regions of interest in the TONIC cohort. The colored squares indicate clusters of features that are characteristic of a distinct biological process, e.g., immune diversity, morphology, or hypoxia. Nuc: nuclear. Cyto: cytoplasmic. Perim: perimeter.

that capture changes at both broad lineage levels and more granular subsets (e.g., total T cells vs. CD8 T cells), diversity scores for assessing the relative balance between multiple populations, and mixing metrics that quantify the degree of partitioning between distinct cell populations (**Fig. 2a**). We used SpaceCat to automatically extract and quantify each of these features in each of the 678 tissue cores included in the dataset.

To systematically capture the influence of location on cells in the TME, we defined four tumor compartments in each image: the cancer core (regions with high cancer cell density), cancer border (the outer edge of these regions), stroma border (the surrounding non-cancer edge), and stroma core (the remaining image area) (**Extended Data Fig. 7a**, Methods). The features described above were computed across the whole image, as well as within each of these compartments to obtain compartment-specific estimates. Some features, such as the proportion of cancer cells positive for Ki67 and the proportion of CD68 Macrophages positive for PD-L1, were consistent across compartments (**Extended Data Fig. 8a, f-g**). In contrast, the ratio of CD8^+^ T cells to CD4^+^ T cells was significantly higher in the cancer core, supporting the hypothesis that CD8^+^ T cells specifically migrate to areas of high cancer density to execute their cytotoxic function^39,40^. In total, 28% of features varied by compartment (**Extended Data Fig. 8c**), predominantly driven by differences between tumor and stromal areas (**Extended Data Fig. 8b**), highlighting the importance of accounting for spatial context.

Of the 872 features calculated by SpaceCat, the most common were those capturing functional marker expression (n=347, i.e. proportion of Ki67^+^ cancer cells) and cellular densities (n=232, i.e. T cell density), which is a reflection of both how informative those features are (since highly correlated features are removed) and the specific markers included in our panel (**Fig. 2b**, **Extended Data Fig. 2a**). Other categories of features included cellular morphology (n=96), spatial mixing between cell types (n=6), cellular diversity (n=38), regional diversity (n=20), cell-cell distances (n=24), and features quantifying the extracellular matrix (n=78). Due to the immunological focus of our antibody panel, we could define immune cells with the highest degree of granularity, and hence they were the most represented cell type in SpaceCat features (42.3%). A correlation matrix of the distinct features revealed clusters of biologically-related features (**Fig. 2c**). For instance, features associated with immune diversity, such as CD4^+^ T neighborhoods, macrophage proportions, and global cellular diversity were highly correlated. We also identified narrower modules, including those related to hypoxia (GLUT1^+^ positivity) and cell morphology (nuclear to cytoplasm ratio and membrane concavities) (**Fig. 2c**). By defining and quantifying the core components of the TME, SpaceCat provides a means to query multiplexed imaging data, generating an interpretable set of features that can be used to separate patient populations from one another.

### Predictors of response to immune checkpoint blockade

To understand how TME structure relates to ICI benefit, we tested the association of each feature from SpaceCat with patient outcome, identifying features that were associated with patients who did or did not respond to nivolumab in the TONIC trial (**Fig. 3a**, Methods). To quantify the strength of these associations, we created an importance score that incorporated statistical significance and effect size, with 0 indicating the least predictive and 1 indicating the most predictive feature. For the subsequent analyses, we focused on the 100 highest-ranking features (**Extended Data Fig. 9**, Methods).

**Figure 3.**
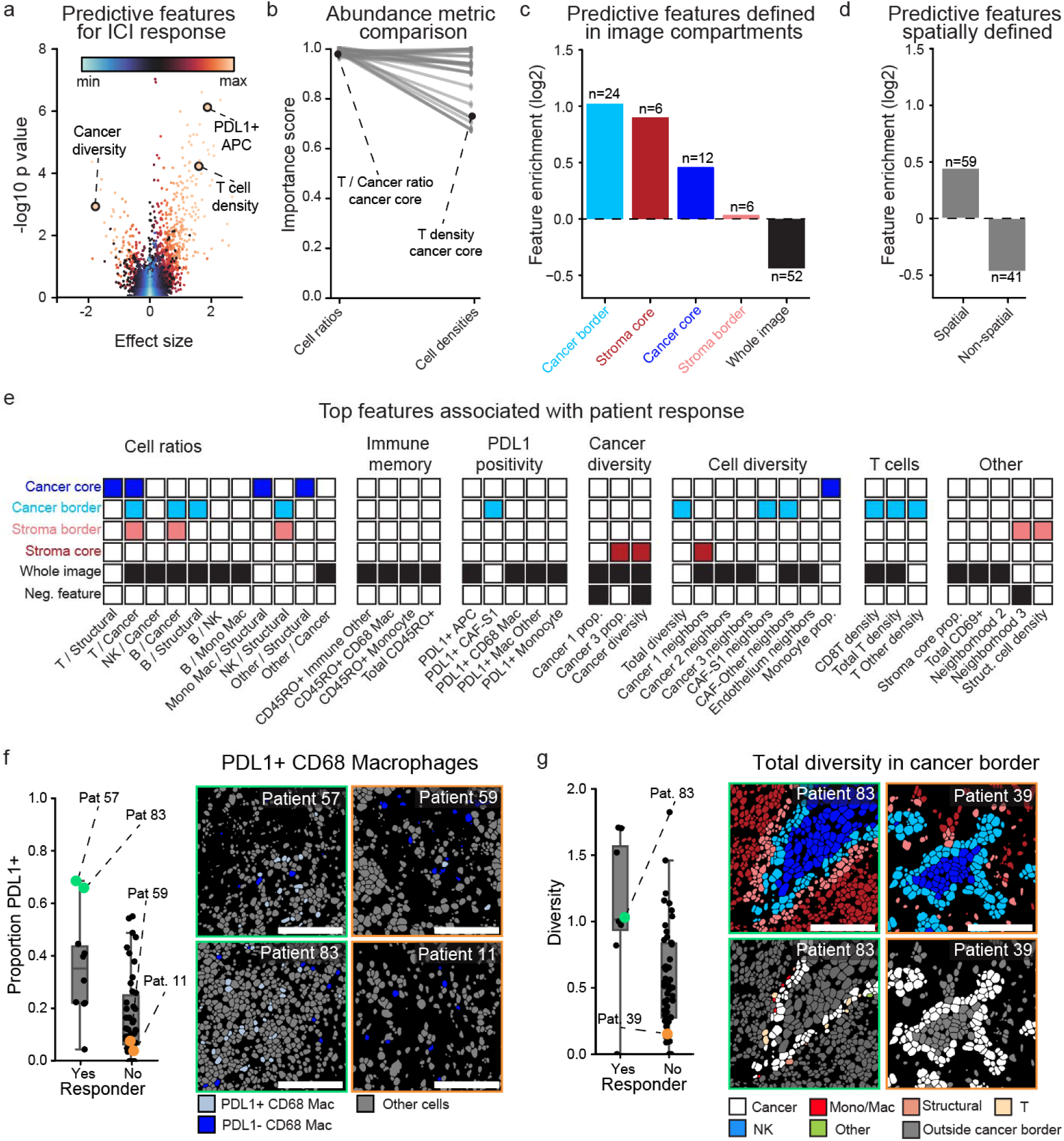
Extracted microenvironmental characteristics associated with patient response. **a**) Volcano plot showing significance (t-test, y axis) and effect size (difference in medians, x-axis) of features to predict patient response, colored by overall ranking. **b**) Comparison of cell ratios and individual cell densities to predict outcome. The score for each of the ratios in the top 100 features is shown on the left hand side. For each ratio, which is composed of two different cell types, the cell type density with the higher score is plotted on the right. T / Cancer ratio; ratio between T cells and cancer cells. T density; density of T cells. **c**) Enrichment within the top 100 features for features calculated within each of the tumor compartments, or those calculated across the whole image. **d**) Enrichment within the top 100 features for features that do and do not require spatial information to be calculated. **e**) Top 50 features associated with response grouped by category. The first five rows for each feature indicate which compartment(s) show(s) an association with outcome for that feature, while the bottom row illustrates whether the feature is positively or negatively associated with outcome. Ratios between cell types are detonated with a ‘/’, e.g. T cell to Cancer cell ratio is T / Cancer. Neighborhood diversity is a spatial metric that takes into account the immediate neighbors of a given cell type, whereas other diversity metrics are calculated using the total count of cells within a compartment. Scale bars: 100um. **f**) Representative example of a top feature (PD-L1+CD68+ Macrophages). The boxplot on the left shows the feature stratified by outcome. The overlays on the right show four specific examples (highlighted in the box plot) of patients with high and low levels of PD-L1+CD68+ Macrophages. Data is from the on-nivo timepoint. **g**) Representative example of a top feature (diversity in the cancer border region). The boxplot on the left plots this feature stratified by responder/non-responder stat s. The overlays on the right show two specific examples (highlighted in the box plot) of patients with high and low border diversity. The top row shows the image compartments (same coloring as in 3c and 3e), and the bottom row shows the cell types present in the cancer border compartment. Data is from the on-nivo timepoint. Scale bars: 100um.

We first evaluated if specific feature classes from SpaceCat were more predictive of response. SpaceCat calculates not only densities for individual cell populations, but also how the density of one cell population varies relative to another (e.g. the ratio of macrophages to T cells). When comparing these distinct abundance metrics, we found that cell ratios occurred more frequently in the top 100 features compared to cell densities (**Extended Data Fig. 10a**). We compared the importance score of each ratio in the top 100 features with the higher-ranked of the two cell densities used to calculate it. We found that ratio-based scoring consistently outranked the individual cell densities, suggesting that the additional information captured in the cell ratio drives increased association with outcome (p<0.0001, **Fig. 3b**). For example, the ratio of T cells to cancer cells in the cancer core compartment had an importance score of 0.99, whereas T cell density had a score of 0.71, and cancer cell density had a score of 0.62.

Next, we sought to understand the importance of spatial information for predicting outcomes. We observed substantial enrichment within the top 100 most predictive features for those defined within a specific tumor compartment (p<0.0001, **Fig. 3c**). For example, the global diversity of cells within the cancer border was the 31st highest ranked feature, while the same metric evaluated across the entire image ranked 302nd. Taken as a whole, spatial features were significantly enriched within the top 100 features compared to non-spatial features (p=0.001, **Fig. 3d**).

We organized the top-ranking SpaceCat features into modules based on their underlying biological processes (**Fig. 3e**). Looking at the ratio-based features, we saw increased relative abundance of lymphocytes to cancer or structural cells to be consistently associated with better outcome (e.g. larger values of the T cell / cancer cell ratio, B cell / cancer cell ratio, and T cell / structural cell ratio were beneficial; **Fig. 3e**, “Cell ratios”). In line with prior work^37,41^, another module consisted of PD-L1 expression on non-cancer cells (**Fig. 3e**, “PD-L1 positivity”). For example, the proportion of PD-L1+ CD68^+^ macrophages was significantly higher in responders compared to non-responders (p=0.002, **Fig. 3f**).

Looking at both neighbor-based and abundance-based diversity metrics (**Extended Data Fig. 7**), we found that increases in the diversity of the overall composition of the images were consistently associated with better outcomes (**Fig. 3e**, “Cell diversity”). This included the diversity of cells within specific image compartments (**Fig. 3g**), diversity surrounding cancer and structural cells, and diversity within monocytes. In contrast, diversity specifically within cancer cells was not exclusively associated with better outcomes. A higher proportion of the Cancer 1 population (defined by expression of epithelial markers ECAD and CK17) was negatively associated with outcome, whereas a higher proportion of Cancer 3 (defined by low expression of epithelial markers) was positively associated with outcome. Total cancer diversity (encompassing Cancer 1, Cancer 2, and Cancer 3) was negatively associated with patient outcome (**Fig. 3e**, “Cancer diversity”).

To understand the generalizability of these findings, we analyzed the bulk RNA-seq data from the same samples. We applied 134 previously defined RNA-seq signatures to our data to summarize the key components of the TME (Methods). We then used the same approach as above to generate importance scores summarizing the degree to which each RNA-seq signature predicted response (**Extended Data Fig. 10b**). Looking at the top features, we observed two broad categories. The first reflected increased immune cell infiltration, such as signatures of effector cells and B cell infiltration (**Extended Data Fig. 10c**), in agreement with the MIBI-based features of cell diversity and T cell abundance. The second category captured aspects of immune cell state such as cytokine secretion and interferon signaling that were not well-covered by our MIBI panel (**Extended Data Fig. 10c**), demonstrating the potential of RNA-seq data to complement our MIBI analysis.

### Response associated features evolve through time

One of the major strengths of our cohort is longitudinal sampling of the primary, baseline, pre-nivo, and on-nivo samples from the same patients. The preceding analyses generated a single timepoint-agnostic ranking to give a global view of the TME features associated with ICI response. With this foundation in place, we next sought to understand to what extent these properties were temporally conserved. This revealed two insights. First, 92 of the top 100 features were associated with outcome at only one timepoint (**Extended Data Fig. 10f**). Second, the predictive value of the information gleaned from each timepoint was not equivalent, with the majority (n=80) coming from the on-nivo samples (**Fig. 4a,b**, **Extended Data Fig. 10g**).

**Figure 4.**
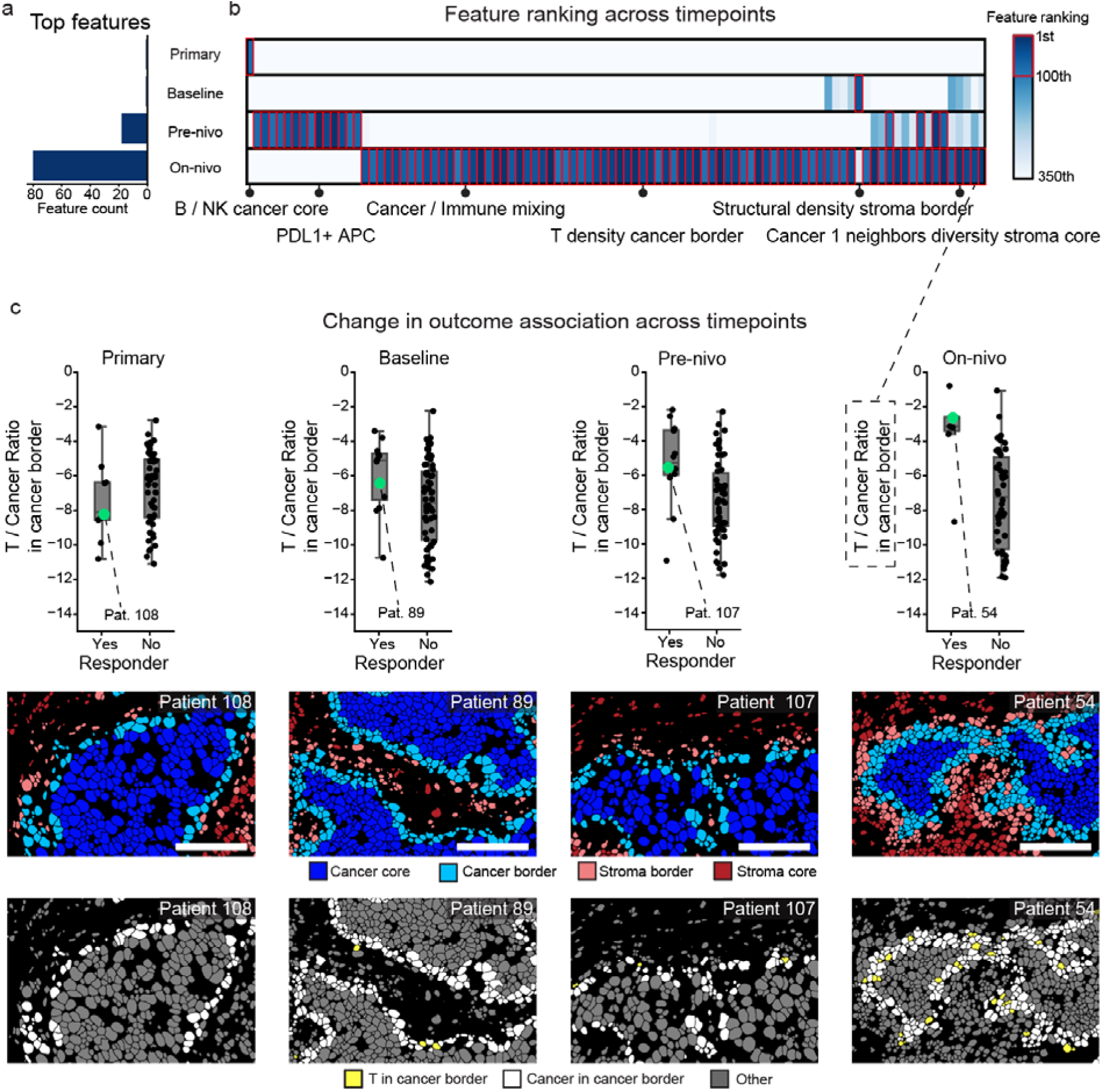
Evolution of features associated with response. **a)** The number of features from the top 100 that are derived from each timepoint. **b**) Heatmap showing the overlap of the top features across different timepoints. In order to be included in the visualization, a feature needs to be within the top 100 most predictive. Using this list of features, we then plot those features in any timepoint where a feature is ranked within the top 350 features. Features are colored by their overall ranking, and boxed in red if they are within the top 100. **c**) The top row of box plots shows the ratio of T cells to cancer cells within the cancer border broken down by response. This is plotted across all four timepoints to show the change in association with outcome. Underneath are representative overlays showing the compartments within each image, followed by the T cells and cancer cells within the border compartment. Scale bars: 100um.

This prompted us to look in more detail at the top features from each timepoint. The top three features from the primary tumors were the B cell / NK cell ratio in the cancer core, the density of CD68 macrophages, and the proportion of CD4^+^ T cells positive for Vimentin (**Extended Data Fig. 11a-c**). Even though these features were top ranked in the primary tumor samples, the degree of separation between responders and non-responders was modest, and none were significant at other timepoints. Features that are typically referenced as prognostic in primary TNBC^38,40^, including T cell / cancer cell ratio, PD-L1^+^ APCs, PD-1^+^ CD8^+^ T cells and T cells close to tumor cell, did not predict subsequent response to ICI when assessed in the primary tumor samples and ranked surprisingly low (below 1000).

We next looked at the baseline samples to identify the types of features unique to this timepoint. Overall, the most represented feature categories at baseline (**Extended Data Fig. 10i**) were cell ratios and cell diversity. The top three features at baseline were the density of structural cells in the stroma border, the ratio of unclassified (other) cells to cancer cells in the stroma core, and the ratio of NK cells to other cells in the stroma border (**Extended Data Fig. 11d-f**). None of these features were predictive in other timepoints, highlighting the timepoint-specific relationship these features have with response.

The on-nivo timepoint had the largest number of distinct features associated with outcome, as 92 out of the top 100 features originated from this timepoint, many of which related to T and B cell densities, as well as ratios between T and B cells and other cell types (**Extended Data Fig. 10g**). In contrast, the baseline features did not relate to T and B cells to nearly the same degree, as the most important baseline features were related to cancer and structural cell diversity (**Extended Data Fig. 10g,h**). Top hits from the on-nivo tumors included features aligned with previously published findings – such as the ratio of T cells and B cells vs. cancer cells^40^, the expression of PD-L1 on macrophages^20^, and the expression of CD69 as a marker of tissue residence^39^ – in addition to other correlates of outcome, such as cancer cell and fibroblast diversity. Close physical proximity between cancer cells and fibroblasts was positively associated with patient outcome, suggesting that these interactions may play a role in driving good outcomes. One of the highest ranked features was the ratio between T and cancer cells in the cancer border compartment. Notably, no significant association between this feature and outcome was observed in the primary, baseline, or pre-nivo samples, consistent with an influx of T cells to the border region being an early indicator following initiation of ICI for patients who will go on to respond (**Fig. 4c**).

Although most features were unique to one timepoint, four features in the top 100 were shared across timepoints (**Fig. 4b**, **Extended Data Fig. 10f**). Of these, three out of four (75%) were specific to the cancer border region, including cellular diversity, PDL1^+^ CAF-S1 cells, and the ratio of other cells to cancer cells. We next assessed how the degree of overlap increased when using a more permissive threshold of the top 350 features (**Fig. 4b**). Looking across the top 350, we identified 24 features shared across at least two timepoints and six features that were shared across three timepoints. All six of these features were shared across the metastatic timepoints, and none overlapped with the primary tumor. Two of these features were specific to cancer border, including neighbors diversity of CAF-Other cells and the T cell / cancer cell ratio (**Fig. 4c**). Thus, we found a strong temporal dependence of the identified features, with the majority coming from the on-nivo timepoint. Some features predicted response across baseline, pre-nivo, and on-nivo tumors, but the majority were unique to only a single timepoint, corresponding to significant shifts over time in the specific features which predicted response.

### Multivariate modeling to predict patient response

Having identified multiple features that were individually associated with response to ICI, we next sought to develop a multivariate model to predict treatment response at each timepoint. In brief, we performed classification using a Lasso model to predict whether a patient was classified as a responder or non-responder. We used all 872 SpaceCat features to train a separate Lasso^42^ model on each timepoint, using nested cross validation to generate estimates of the accuracy of each model (**Extended Data Fig. 12a**, Methods).

A model trained exclusively on MIBI data from the primary tumor had poor performance (mean AUC=0.62), which was modestly better than random chance (**Fig. 5a**). In contrast, we observed substantially better performance for the models trained on the baseline (p<0.0001, mean AUC=0.77) and pre-nivo tumors (p=0.003, mean AUC=0.69). To contextualize these AUC values, we compared our results with those of a recently published analysis of the NeoTRIP^20^ trial, which evaluated the efficacy of neo-adjuvant immunotherapy for early-stage TNBC using multiplexed imaging. A similar AUC value for predicting response to immunotherapy using pre-treatment biopsies (0.77) was reported in this study compared to our AUC for the model trained on the baseline metastatic samples (0.76). Interestingly, identical accuracy for the on-treatment samples was reported in that study, which also had an AUC of 0.77. In contrast, we observed substantially higher accuracy from the on-treatment timepoint in TONIC, with an average AUC of 0.91 (**Fig. 5a**). The relative ranking of the timepoint-specific models is in line with our findings from the univariate analysis (**Fig. 4b**), where numerous features from the on-nivo timepoint were associated with outcome and almost no features from the primary timepoint were associated with outcome.

We used the same approach to evaluate timepoint-specific models on the bulk RNA-seq data. Looking across timepoints, we observed a similar overall trend, with the RNA model trained on the on-nivo timepoint (AUC=0.88) outperforming the RNA models trained on the pre-nivo (p<0.0001, AUC=0.56) and baseline (p<0.0001, AUC=0.73) timepoints. Interestingly, both RNA and MIBI models achieved excellent performance at the on-nivo timepoint, while the MIBI model for the pre-nivo (p=0.003) and baseline (p=0.06) timepoints outperformed the corresponding RNA models. Thus, for samples taken before ICI treatment is initiated, the MIBI-based models performed the best. Unsurprisingly, a model trained on baseline genomic features (somatic mutations and copy number alterations) was not as accurate at predicting outcome (AUC=0.67) (**Fig. 5a**). Overall, this highlights the importance of longitudinal sampling and the value of early on-treatment biopsies, consistent with our prior findings in the context of neo-adjuvant anti-HER2 targeted therapy in early-stage breast cancer^43^.

**Figure 5.**
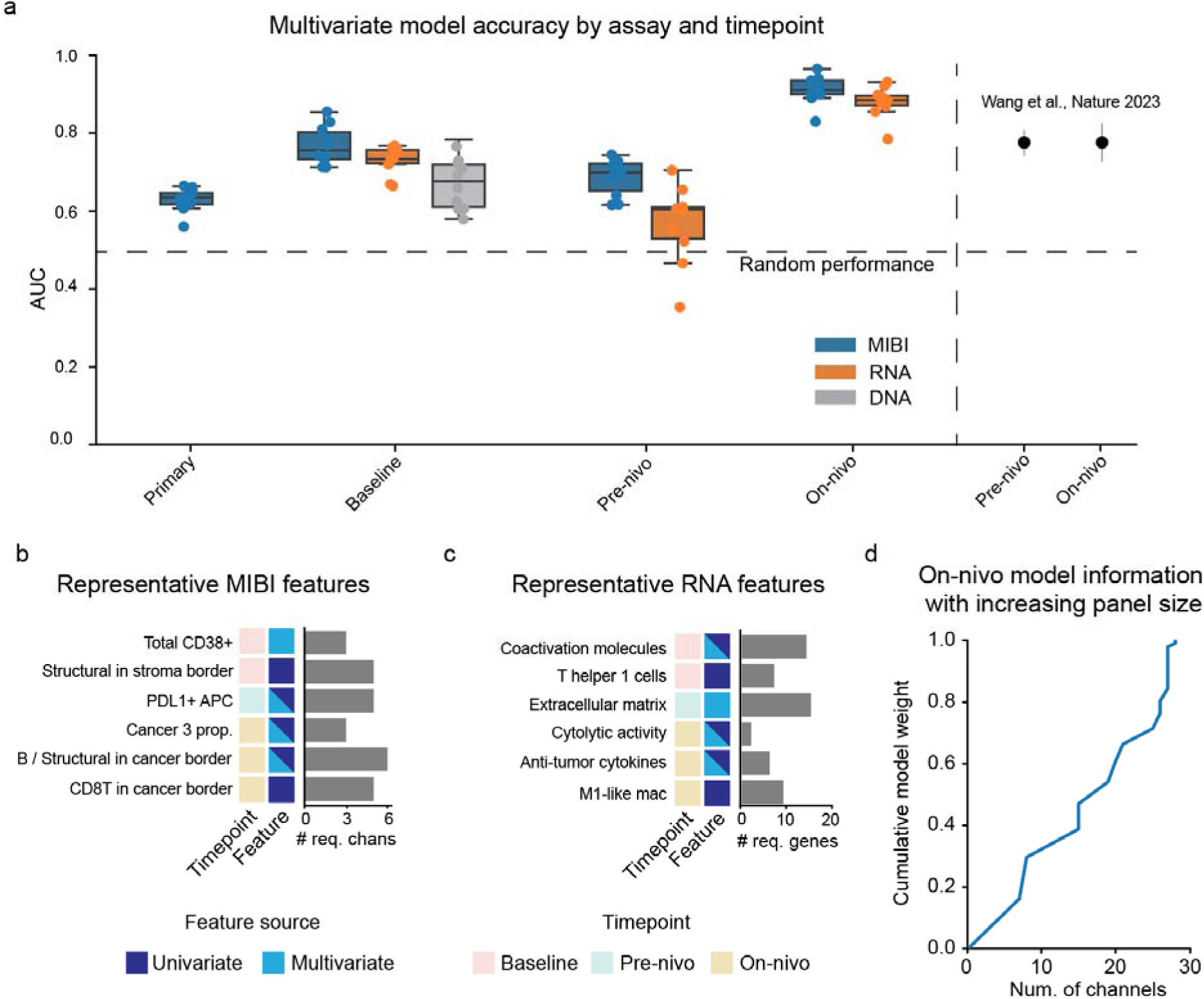
Multivariate modeling to predict response. **a**) The AUC (y axis) of the multivariate models stratified by assay and timepoint (x axis). Each dot is a replicate from nested cross validation. On the right, data from Wang et al. 2023 was replotted on the same axis. **b**) Representative features derived from the MIBI data that were strongly associated with patient outcome. The number of distinct channels required to calculate each feature is shown with a horizontal bar, along with the relevant timepoint and the analysis method (univariate or multivariate) that identified the feature. **c**) Same as b), but for RNA-based features, showing the number of transcripts instead of number of channels. **d**) The cumulative sum of the weights of the features that can be calculated (y axis) as more channels are included in the imaging panel (x axis).

One of the strengths of using Lasso is its interpretability, as each feature is assigned a linear coefficient that is proportional to its importance. We used these coefficients to examine the specific features that were selected by each model (**Fig. 5b-c**) and the data requirements for generating them. Interestingly, the baseline model identified a number of features as important predictors, including CD38^+^ cells (**Fig. 5b**). CD38 is a glycoprotein with roles as both a receptor and an enzyme; recent studies have indicated that CD38 may play an immunosuppressive role in the TME during ICI^44,45^. Of note, we found CD38 to be mostly expressed by endothelial cells, whereas previous work found T cell and cancer cell expression to be informative. CD38 positivity requires only three markers to be evaluated (two for segmentation, plus CD38), and even fewer if manual scoring is used, facilitating clinical implementation. The top feature for the baseline RNA model was the cytolytic activity score, a two-gene score proposed by Rooney et al. that averages the expression of two cytotoxic granule genes, Perforin-1 and Granzyme-A^46^.

The on-nivo MIBI model identified nine features as important predictors, including cancer diversity and the ratio of B cells to structural cells. The identified MIBI features would each require between three and six distinct markers to be computed (**Fig. 5b**). The RNA model identified three important features: the cytolytic activity score, a signature of anti-tumor cytokines, and an extracellular matrix score (**Fig. 5c**). The cytolytic activity score was also identified by the baseline RNA model, and was shared across all three timepoints (**Extended Data Fig. 12f**). The RNA-based signatures required an average of eight distinct genes for calculation. Finally, we analyzed the relationship between the size of the antibody panel and the number of features the model relies on (**Fig 5d**). For the highly predictive on-nivo model, 28 antibody channels were needed to generate all selected features, out of a total of 37. However, the relatively minor contribution of certain features suggests this number could be reduced, with implications for how our findings could be translated into a scalable assay.

## Discussion

Here, we examined how the TME evolves over time in patients who developed metastatic disease and how these dynamics relate to response to ICI. We constructed a unique clinical cohort spanning primary diagnosis to on-treatment metastatic disease from patients enrolled in the TONIC trial. Moreover, we developed an interpretable, open-source computational framework (SpaceCat) to extract over 800 features from multiplexed imaging data. Looking across primary tumors, pre-treatment metastases, and on-treatment metastases, we performed in-depth spatial, transcriptomic, and genomic characterization to define the core components of the TME. We then assessed to what extent these components were related to ICI response, finding numerous features that were associated with patient outcome. Looking across time, we found the primary tumor to be a poor predictor of patient outcome, whereas the metastatic tumors harbored much more information about the determinants of response. We identified predictive features that were shared across timepoints as well as features that were unique to a specific timepoint, highlighting the importance of collecting longitudinal samples to understand the relationship between the TME and response to ICI.

Our analysis revealed significant insights into the biology underpinning patient responses to ICI. We found strong and consistent associations between cellular diversity and response to ICI. In addition to overall diversity, we also found increased ratios of immune cells to cancer cells to be a robust predictor of patient response, both at the image-wide level as well as at the interface between stromal and cancer compartments^47–49^. Moving beyond cell abundance, PD-L1 positivity on APCs, CD68^+^ macrophages, CD163^+^ macrophages, and CAFs in the pre-treatment tumors was predictive of response to ICI. The breadth of features we found to be associated with outcome emphasizes the crucial role myeloid and structural cells play in immune activation and the regulation of the TME.

Surprisingly, we found few predictive features in the pre-treatment tumors that were based on lymphoid cell function or abundance. For instance, pre-treatment density of T cells did not robustly stratify patients, nor did T cell expression of previously identified exhaustion markers. In contrast, the on-treatment tumors displayed numerous features related to T cell localization and abundance that predicted ICI response. This suggests that T cells in the TME may not be the only mediators of ICI response and that these therapies are exerting their influence by modulating the immune system at multiple levels^50^. This observation aligns with recent research suggesting that the infiltration and replenishment of new immune clones, rather than the expansion of existing ones, is the predominant pattern in responsive patients^47–49,51,52^.

We observed a significant increase in both the number and strength of features associated with patient response at the on-treatment timepoint compared to the pre-treatment timepoints. Although it might seem intuitive that an on-treatment timepoint would exhibit the most pronounced signs of response, two recent studies of early TNBC where patients received neoadjuvant ICI did not observe more informative on-treatment features compared to pretreatment features. In the NeoTRIP trial^20^, the authors used multivariate models to predict patient outcome and observed equal accuracy from models trained on the pre-treatment and on-treatment samples. Similarly, Shiao et al. identified two distinct response trajectories for patients responding to ICI but did not find an enrichment of these features at the on-treatment timepoint^53^. It is important to note that the patients in the TONIC trial had relapsed metastatic disease and had undergone multiple lines of prior therapy, unlike the untreated primary disease cohorts in the aforementioned studies. Moreover, the patients receiving neoadjuvant ICI for primary disease also received concurrent cytotoxic chemotherapy. Thus, it could be that chemotherapy treatment in these patients masked the impacts of ICI on the TME. The differences in the information content of on-treatment samples in our study compared to prior work are thus likely a combination of different biology in the metastatic setting, coupled with different patient populations.

We found that the antecedent primary tumor had relatively little ability to predict subsequent ICI response for the metastatic patients enrolled in the TONIC trial. This could be due to the evolution that the metastases have undergone in response to multiple lines of treatment or the metastatic process itself. An important implication of this finding is that studying the determinants of treatment response in the metastatic setting is likely to benefit from the collection of metastatic lesions, ideally as close as possible to the start of treatment. Despite the simplicity of this statement, there are numerous practical, logistical, and ethical considerations that have prevented it from becoming the norm. Significant time and resources will need to be dedicated to enable the collection of such samples, along with dialogue with patient advocates and clinicians to underscore why these samples are so crucial. Considering that response monitoring in the clinic is becoming more adaptive and patient specific^54^, it is likely that in the future, patients will be routinely monitored with serial imaging and liquid biopsies to inform their treatment. This is already being tested in several clinical trials^55–59^ to help clinicians and patients make informed decisions about treatment efficacy and timing.

The multi-modal nature of our dataset enabled us to compare information from MIBI, bulk RNA sequencing, and bulk exome sequencing of the same samples. This raises an important question: What data is essential for a better understanding of the TME? Our results indicate that studies aimed at quantifying and characterizing the TME would benefit most from modalities that capture more than just alterations to patients’ DNA. In the TONIC trial, we did not find strong correlates of response at the DNA level. Specifically, driver mutations, copy number alterations, and the degree of genomic disruption did not reliably predict ICI benefit in metastatic TNBC patients. Although multivariate models trained on the RNA-seq data from the on-treatment timepoint were just as accurate as those trained on the MIBI data, the MIBI models were more accurate at the pre-treatment timepoints. In addition, the RNA-seq features were arguably less informative. For example, knowing that the density of T cells in the border region of a tumor was a top predictive feature provides more fertile grounds for hypothesis generation than knowing that “Effector cells” was a top RNA signature associated with response. The difficulty of deconvoluting bulk RNA-seq data into signatures that truly represent individual cell populations adds a further challenge when interpreting specific signatures. In our work, we found that MIBI and RNA-seq were equivalent in their ability to predict response, but that the MIBI data provided more actionable features for subsequent follow-up.

Our study has several limitations to consider. Foremost, the TONIC trial was undertaken specifically to profile patients with metastatic TNBC. Thus, it is not clear to what extent our findings here will generalize across breast cancer subtypes, or to other cancer types. Due to the scarcity of longitudinal samples in metastatic ICI trials, we could not validate our findings in an independent cohort that mirrored the underlying characteristics of our patient population. To make the generation and analysis of imaging data feasible, we constructed tissue microarrays using pathologist-identified regions from each patient’s tumor, rather than analyzing the entire tumor block. This approach may miss informative features for these patients. Overall, our study underscores both the importance of spatial modalities for generating accurate and informative TME features to correlate with and predict patient outcome on ICI, as well the potential translatability of more easily implemented assays like bulk transcriptomics to deliver similar predictive power, though at the cost of interpretability.

## Methods

### Study design and sample collection

The TONIC trial (NCT02499367) is an adaptive phase II randomized, non-comparative study evaluating the feasibility and efficacy of nivolumab following a 2-week induction treatment in patients with metastatic TNBC. This single center, non-blinded trial was conducted in two stages, according to Simon’s two-stage design^60^. Initially, five cohorts were included; four receiving induction treatment (low-dose chemotherapy or irradiation) before nivolumab and one without induction treatment. Results of the first stage of the trial were previously reported^24^. In the second stage, the number of arms was reduced based on the first stage results, following the ‘pick-the-winner’ approach and considering both clinical and translational endpoints. For this translational study, patients from both stages I and II were included if they had at least one tissue sample available (n=123 out of 127 eligible patients, clinical data published in^24,25^).

All patients biopsies from metastatic lesions were taken at baseline of the TONIC trial (baseline). Core biopsies from metastatic lesions were taken before the start of the study immunotherapy (baseline), after induction and after three cycles of nivolumab (240 mg flat dose) ^24^. To explore the additional value of microenvironmental analyses from the primary tumor, we retrospectively pursued to collect the paraffin-embedded archival tissue blocks from each patient’s therapy-naive primary tumor **(Fig. 1a)**. For all patients accrued up to 2019/07/23, we requested the archival formalin-fixed paraffin-embedded (FFPE) tissue blocks of the primary, therapy-naive, tumor. We received representative tissue blocks of the primary tumor for 117 patients. Archival tissue blocks were manually screened by a breast cancer pathologist in slidescore^61^. New hematoxylin and eosin–stained (H&E) whole slides were prepared for all tissue specimens and histopathologic features were reexamined by a dedicated breast pathologist. Each H&E section was analyzed to determine which samples contained tumor cells, and were most representative of the tumor (**Extended Data Fig. 3**). Then, regions of tumor that contained stroma were annotated, cored, and assembled into 21 tissue microarrays (TMAs) of 1.5□mm cores.

Key inclusion criteria were: ≥18□years; metastatic or incurable locally advanced TNBC with confirmed estrogen receptor negativity (<□10%) and HER2 negativity (0, 1+ or 2+ without amplification determined by in situ hybridization) on a biopsy of a metastatic lesion or breast recurrence. Additional criteria were detailed previously^24^. Neoadjuvant chemotherapy was given to 55.6% of patients for their primary tumor, with only minority achieving a near-complete (3.4%) or complete (6.8%) pathological response at surgical resection, consistent with poor prognostic outcomes^62–64^. Adjuvant chemotherapy was given to 38.4% of patients. Responders were defined by a best overall response of complete response (CR), or partial response (PR) according to RECIST1.1^65^ and iRECIST^66^. The trial was conducted in accordance with the protocol, Good Clinical Practice standards and the Declaration of Helsinki. The full protocol and the informed consent form were approved by the institution’s medical-ethical committee. All patients provided written informed consent before enrollment.

### Control TMA construction

We constructed a control tissue microarray to identify slide-to-slide variation in staining. Each core on this TMA was 1.5mm, for a total of 13 cores. Control tissues were carefully selected from archival FFPE tissues from the Stanford pathology department. The control TMA included two replicate cores each of tonsil, spleen, lymph node, breast (DCIS, IDC, and normal areas), colon and placenta, plus an additional tonsil for asymmetry. Serial recuts from the control TMA, along with from the cohort TMAs, were cut onto the same slide. Both TMAs on each slide went through all subsequent processing steps in parallel.

### MIBI staining

#### Panel construction

The majority of the antibodies in this study have been previously validated for MIBI^37,67,68^. New target antibodies were first validated by immunohistochemistry to confirm appropriate staining patterns in control tissue samples. All antibodies were then metal-labeled with the Ionpath conjugation kit (IonPath, Menlo Park, USA) following the manufacturer instructions. To increase reagent shelf life, labeled antibodies were then lyophilized individually with 100 mM trehalose in aliquots of 1 ug or 5 ug format. Following lyophilization, the appropriate antibody titer was determined by serial dilution with the following starting titer range (1 ug/mL, 0.5 ug/mL, 0.25 ug/mL, 0.125 ug/mL), for the new targets, or with the recommended titer, for the relabeled MIBI validated antibodies.

#### Cohort staining

To reduce batch effects, all TMAs were stained with the same mastermix. Fresh aliquots of each antibody were reconstituted and combined together into a single mastermix. Each step in the staining protocol was performed in pairs, with one reader and one pipettor, to reduce mistakes. For a complete description of the experimental conditions and procedures used, see our methods publication^69^. For a step-by-step guide, see below.

#### Interactive protocols

Reagent preparation: https://www.protocols.io/view/mibi-and-ihc-solutions-261geo7wyl47/v1

IHC staining: https://www.protocols.io/view/ihc-staining-x54v9moxmg3e/v1

MIBI staining: https://www.protocols.io/view/mibi-staining-dm6gprk2dvzp/v5

Sequenza staining: https://www.protocols.io/view/staining-sequenza-6qpvrdeo2gmk/v1

Antibody lyophilization: https://www.protocols.io/view/antibody-lyophilization-kxygxex5kv8j/v1

### MIBI data generation

#### MIBI run setup

All MIBI data was generated on a commercial MIBIScope instrument (IonPath, Menlo Park, USA). For each TMA core on each TMA, we used paired H&E images to identify the subregion that would be acquired on MIBI, avoiding areas of necrosis or empty slide (**Extended Data Fig. 1b**). TMA cores were named by their row and column number in the TMA grid to enable easy mapping back to the appropriate metadata. To ensure that the cores were reliably assigned the correct row and column coordinates on the TMA, we added automated checks to identify naming errors and fix them following user confirmation. Prior to beginning data acquisition, the order of the cores was randomized to mitigate potential batch effects from instrument drift.

#### Acquisition settings

We used the same settings across all MIBI data acquired in this study. The size of each field of view (FOV) was determined based on sample availability. When possible, FOVs of size (800 um)^2^ and (2048 pixels)^2^ were acquired. However, if insufficient tissue was present, an FOV size of (400 um)^2^ and (1024 pixels)^2^ was used instead. We used a custom preset of 8.0 nano Amp beam current and 0.63 millisecond dwell time to balance acquisition speed and image clarity. We disabled all default background correction and noise removal settings.

#### MIBI cohort features

Multiple regions of interest (ROIs) per tissue sample were selected based on the pathologist annotation for areas representative of the tumor and stroma. A total of 1614 cores from 123 unique patients were included on the TMAs for MIBI data acquisition **(Table S1)**. MIBI data was successfully generated for 1256 cores (success rate 78%), and includes cores from 117 unique patients. We used MIBI to generate high quality, multiplexed imaging data with subcellular resolution across all 21 TMAs in our study. The total dataset consists of 1256 fields of view (FOV), 678 were included in this project.

### MIBI data processing

#### Image compensation with Rosetta

To systematically correct the sources of contamination and background present in the MIBI data, we used an approach analogous to flow-cytometry channel compensation called Rosetta. To do this, we linked sources of potential contamination or spillover, such as spillover from the gold channel, organics, and adducts (source channels) to the channels where that signal showed up (target channels). We used previously published estimates^70^ as well as updated values from the MIBIScope manufacturer (IonPath, Menlo Park, USA) to derive coefficients for isotopic impurities in the metal conjugates, as well as instrument-specific coefficients for elemental background in each channel (e.g. adducts and oxides). To account for organic signal, we extracted a range of the spectrum (AMU 117-125) without any true signal to serve as a template for non-specific signal, colloquially referred to as “noodle” due to its characteristic appearance. We also used the gold channel as a proxy for slide background, since the slides are gold-coated. We manually identified the appropriate coefficients for the gold and noodle channel by iterative refinement.

Following identification of the appropriate coefficients for each source of contamination, we constructed a matrix to hold the coefficients. Each row in the matrix represents a source channel, and each column represents a target channel. The value of each entry *ij* in the matrix represents the sum of all sources of contamination from the mass in row *i* for the target channel in column *j*. This quantifies the proportion of signal in each source channel that appears in the target channel. To correct the target image, we subtracted the source channel multiplied by its coefficient from the target channel (**Extended Data Fig. 3a**). We used gaussian smoothing to convert the integer counts into decimals to enable fractional compensation. Representative images taken before and after noise removal with Rosetta can be seen in **Extended Data Fig. 3b**.

#### Intensity normalization using median pulse height

Over the course of a run, the MIBI instrument will gradually lose sensitivity due to aging of the ion detector. Calculating this decrease in sensitivity by looking directly at the image data is challenging, because it is difficult to tease out whether a given change is due to biological or technical reasons. When an ion hits the detector in the MIBI, it produces an electrical pulse. Pulses over a threshold height are recorded as ion hits, whereas pulses under this threshold are discarded. This produces the count-based data that the user interacts with. Over the course of a run, the height of these pulses decreases, such that ions with the same intensity will record shorter and shorter pulses, with more and more of them falling under the threshold. When the sensitivity of the instrument is adjusted, the voltage to the detector is increased such that the height of these pulses is higher. However, looking at the binarized count data (the number of pulses over the threshold), it is challenging to determine if the decrease in counts is due to a decrease in the height of the pulses (technical, decrease in instrument sensitivity), or a decrease in the number of pulses (biological, less signal in the sample).

To circumvent this issue, we used the median pulse height (MPH) to derive an estimate of the purely technical decrease in sensitivity (**Extended Data Fig. 4a**). Because the height of the pulses is determined exclusively by the voltage supplied to the detector, it is independent of the amount or intensity of the protein staining in a given image. Therefore, by calculating the median of the heights for each channel, we can get a robust, independent estimate of instrument sensitivity. Importantly, we can calculate this quantity directly from the image data being acquired for the study, obviating the need to repeatedly measure control samples over the course of the run.

To use the MPH values to correct for instrument drift, we quantified the relationship between MPH and sensitivity. To do this, we constructed a tuning curve. We used a synthetic polymer, poly-methyl methacrylate (PMMA), sample with fixed ratios of metal isotopes to ensure that any change in signal over was due purely to changes in the instrument setup, not sample-specific differences. We then systematically increased the detector voltage, and hence the MPH, and calculated the change in signal, performing this for three replicates at each detector setting. We normalized the resulting signal by the maximum observed, which allowed us to construct a graph relating MPH to the percentage of maximum signal, which we refer to as sensitivity. We then fit a polynomial to this curve, which we can use to convert MPH values to sensitivity. We use the same sensitivity curve for all images.

After generating the sensitivity curve, we calculate the MPH for each channel in each image in a given run. Because the estimate of MPH can be noisy, we generate a per-mass curve that we fit over the course of a run (**Extended Data Fig. 4b**). We use the fitted value to generate a value for the MPH of each mass in each image. We then plug that MPH value into the sensitivity curve to generate the per-mass, per-image sensitivity (**Extended Data Fig. 4c**). Finally, we use this sensitivity estimate to normalize each image, dividing by the sensitivity to bring the values up to 100% sensitivity. Looking at the pre-normalized images over the course of a run, we can see a decrease in signal, especially in the second half of the run (**Extended Data Fig. 4d**). However, following normalization, this decrease in signal is no longer apparent (**Extended Data Fig. 4e**).

### MIBI data QC

To identify potential batch effects in the data, we measured variation in signal intensity across slides, as well as the spatial variation across cores within each slide. To identify potential batch effects across slides, we used the control samples present on each as a reference. We then computed the distribution of the mean of non-zero pixels in each channel in each control sample. Overall, we saw relatively minor shifts in intensity across slides (**Extended Data Fig. 4f**), and the one channel that showed the strongest differences was Calprotectin, which had very few positive cells in the control samples, and thus noisy estimates of positive signal. To identify if different slide locations displayed different sensitivity, we calculated the same metrics as above across each of the cores from the TMA samples. We did not observe any spatial bias in signal intensity across the slides in the cohort (**Extended Data Fig. 4g**).

### Cell segmentation

We used Mesmer^27^ to segment all images in the cohort. Mesmer is a pre-trained deep learning model that takes two channels of input data, a nuclear marker and a membrane marker. We combined the H3K27me3 and H3K9ac channels to form a single nuclear channel, and the CD14, CD38, CD45, CK17, and ECAD channels to form a single membrane channel (**Extended Data Fig. 5b**). We then ran Mesmer using the default parameters as part of the ark-analysis pipeline^71^. Representative images of the resulting segmentation can be found in **Extended Data Fig. 5c**.

### Cell clustering

#### Pipeline overview

We used Pixie^28^ to cluster all of the cells in the cohort. Pixie is a cell clustering algorithm developed specifically for multiplexed imaging data. The first step is to cluster the individual pixels in each image. This produces more robust and reliable estimates of marker expression within each pixel than just using the raw expression values, as we can take advantage of marker correlation patterns. Using these labeled pixel clusters, we then perform cell clustering based on the number of pixel clusters of each type in each cell, rather than using the raw marker values. Following generation of the labeled cell clusters, we manually examined representative images to identify errors in the clustering, which was repeated as necessary. Finally, we performed post-clustering cleanup to address any remaining issues. We used the ark-analysis^71^ pipeline to run all of the described clustering and cleanup steps.

#### Pixie clustering

We performed pixel clustering using the lineage markers contained in the MIBI panel (**Fig. 1f**). We overclustered the data into 225 distinct pixel clusters, which were then combined into 33 meta clusters. We labeled these meta clusters based on marker co-expression patterns, such as CD3+/CD4+/CD45+, ECAD+/CK+, etc, as described in the original publication. We modified the pipeline slightly to improve the clustering on our data. To ensure that channels with significantly higher intensity values did not swamp the per-pixel signal, we added a 99.9th percentile normalization step prior to pixel clustering. To reduce the impacts on downstream cell clustering from noisy channels, we also added the option to exclude non-nuclear signal from nuclear markers (which we used on the FOXP3 channel), as well as the option to remove nuclear signal from non-nuclear markers (which we used on the CD11c channel). We also removed the pixels with the 5th percentile lowest total marker expression, as these were mostly noise and interfered with subsequent cell clustering. Following generation of the pixel clusters, we calculated the proportion of each pixel cluster present in each segmented cell. We then used these proportions as more robust estimates of marker expression in each cell, and fed this into the second step of Pixie, cell clustering. We first over clustered the data into 225 clusters, which were then combined into 33 meta clusters.

#### Cluster inspection and cleanup

To confirm accurate clustering, images were manually inspected using Mantis Viewer^72^. Representative images were selected, and cell assignments were manually verified by toggling on the appropriate image channels, which were then overlaid with the segmentation boundary and cluster assignments. Systematic errors in clustering were then addressed by reclustering either the pixel clusters, cell clusters, or both, as appropriate. Following final assignment of cell clusters with Pixie, an additional round of cleanup was performed using manual thresholds to break up ambiguous or mixed clusters, and to merge duplicate clusters. This resulted in 33 total clusters, which we refer to as ‘detailed’ level clustering because they are the most granular. These 33 detailed clusters were merged into 21 intermediate clusters and 8 broad clusters to facilitate analyses at different levels of granularity (**Extended Data Fig. 6a**).

### SpaceCat

#### SpaceCat overview

SpaceCat computes a wide range of distinct features to comprehensively summarize the tumor microenvironment. This includes features related to functional marker expression, cell density ratios, diversity, mixing proportions, cell neighborhoods, and more. Each image is broken up into four parts (cancer core, cancer border, stroma border, stroma core), and features are calculated at both an image-wide level, as well as separately within each compartment. We set thresholds for the minimum number of cells needed for many of these calculations, and features which do not meet these thresholds in a given image are not computed. We filter out compartment-level features that are highly correlated with the image-level features to reduce redundancy. Following generation of each of these features, they are transformed into a standardized format, Z-scored, and combined into a single data frame for downstream analysis (**Supplementary Table 4**).

#### Defining tumor compartments

We defined four compartments in each image: the cancer core, cancer border, stroma border, and stroma core (**Extended Data Fig. 7a**). We used a smoothed version of the ECAD channel and the segmentations of the cancer cells to define a cancer mask. We binarized this mask, filled in small holes, and kept the regions that were above the size cutoff. We eroded the cancer mask by 50 pixels to define the cancer core, with the eroded regions being defined as the cancer border. We then expanded the cancer mask by 50 pixels, with the expanded region defined as the stroma border. All remaining area was defined as the stroma core.

We performed two steps to clean up these four compartments. We generated a separate mask to represent slide background (i.e. areas with no tissue), which was excluded from the compartment masks. We also generated a separate immune aggregate mask for large clumps of T and/or B cells. These aggregates were removed from the individual compartments to avoid biasing the compartment-wide estimates of cell abundance. With these compartments defined, we assigned each cell to its respective compartment, assigning cells at the border between two compartments based on maximum overlap with the compartment mask. We calculated two sets of features that depended exclusively on the compartment masks: The area of each compartment in each image, as well as all pairwise ratios of compartment areas in each image.

#### Cell abundance features

To quantify cell type abundance, we calculated three categories of features. The first category was densities of individual cells. For each cell type at the broad and intermediate level of clustering (**Extended Data Fig. 6a**), we calculated the density by dividing the number of cells by the area of the region. Densities were computed across the entire tissue area in the image, as well as in each compartment.

The second category was ratios between cell types. We computed all pairwise ratios between cell types at the broad level of clustering, as well as biologically motivated ratios for cells at the intermediate level of clustering (CD8 T/CD4 T, CD4 T/Treg, CD8 T/Treg, CD68 Mac / CD163 Mac). We set a minimum density threshold of 5×10^-7^ cells per square pixel based on manually inspecting the density histograms and looking at the corresponding images. If one of the density values in the ratio was below this threshold, it would be rounded up to the minimum density to avoid issues with division by zero. If both cell densities were below the threshold, the ratio for that image was not calculated. Ratios were computed across the entire image, as well as in each compartment

The third category was cell type proportions. We calculated this feature for cells at the broad level of clustering that were composed of at least two distinct intermediate cell types. For each intermediate cell type in a given broad cell type, we calculated the proportion of the number of broad cells that the intermediate cell type represented. Proportions were calculated across the entire image, as well as in each compartment.

#### Functional marker frequencies

For each cell type at the intermediate clustering level, we calculated the frequencies of positivity for the functional markers on the panel. We manually inspected images to determine an appropriate marker-specific threshold for positivity and applied the same threshold for a given marker across all cell types. For a given cell type, we computed marker frequencies for each marker that was positive in at least 5% of cells of that type, to avoid markers that were not expressed in a given cell type. For each cell type/marker combination, we calculated the proportion of cells above the marker-specific threshold (**Extended Data Fig. 7b**). We set a minimum cell count threshold of at least 5 cells in an image; we did not calculate marker frequencies in images with fewer than the minimum number of cells. Marker frequencies were calculated across the entire image.

#### Cell morphology features

We previously defined a range of morphology metrics to summarize differences in both cell and nuclear shape and size^27^. These metrics were computed across all of the cells in the dataset as part of our segmentation pipeline^71^. Manual inspection revealed that these metrics could reliably pick up differences in cancer cell morphology, but that morphological shifts in the other cell types were too subtle to be effectively captured by these automated metrics. We therefore included cell size for all cell types, and a non-redundant subset of the morphology metrics only for the cancer cells. We computed the average value for each of these metrics for each cell type across the entire image.

#### Cell diversity features

We calculated a range of metrics to capture the diversity of the cells in the image, which fell into two broad categories. The first category was based purely on cell abundances, not physical proximity. We first identified the clustering granularity used to perform the diversity calculation. We did this both at the broad cluster level granularity, as well as at the intermediate clustering granularity within immune, cancer, and structural cell populations. For each of these clustering resolutions, we extracted the proportion of cells of each cell type that was present. We then computed the Shannon index on these proportions (**Extended Data Fig. 7c**). We computed these diversity scores across the entire image, as well as within each compartment.

The second category of diversity feature was those based on physical proximity. For each cell in each image, we computed the number of cells of each cell type within a 50-pixel radius (**Extended Data Fig. 7d**). We then calculated the same Shannon diversity as above, but based on the proportions within the 50-pixel radius. We then calculated the average of this diversity value across the cells at the intermediate clustering resolution. This value was computed across the entire image, as well as within each compartment.

#### Cell-cell distances

For each cell at the broad clustering resolution, we calculated the centroid distance to the nearest cell of every other type^73^ (**Extended Data Fig. 7e**). We then calculated the average across all of the cells in the image, as well as in each compartment in the image. We removed linear distances that were highly correlated with the density of the target cell type to only retain features that represented spatial structure, rather than increased abundance of the target cell.

#### Mixing scores

We calculated all pairwise mixing scores between cell populations at the broad cluster resolution, with some modifications to how it was previously described^37^. For each cell in the selected populations pairs, we computed the number of surrounding cells in a 50-pixel radius. We then calculated the number of heterotypic interactions as the number of cells of the opposite type within the radius, and the number of homotypic interactions as the number of cells of the same type within the radius (**Extended Data Fig. 7f**). We used the ratio of heterotypic to homotypic interactions as the mixing score, which was averaged across all cells in the image.

#### Cell neighborhoods

We used k-means clustering to define cell neighborhoods in each image as previously described^37,67,68^. We computed the relative proportions of cells at the intermediate clustering resolution in a 50-pixel radius surrounding each cell. We used these proportions as inputs to kmeans clustering, selecting 12 total clusters based on the heatmap of cell frequency loadings, as well as manual inspection of the underlying images and neighborhood assignments. We then calculated the proportion of cells belonging to each of the identified cell neighborhoods across the entire image, as well as within each compartment

#### Fiber features

We used our previously described fiber segmentation pipeline^37,67,68^ to segment out individual fiber objects in the images (**Extended Data Fig. 7g**). For each fiber object, we defined key summary statistics such as length, area, elongation, eccentricity, and alignment with neighboring fibers. We averaged each of these metrics across the entire image, as well as within 512×512 tiles, and added these as features.

#### Extracellular matrix (ECM) image clusters

To capture changes in acellular features, we generated image-level clusters based on the expression level of Collagen, Fibronectin, and FAP. We divided the images into tiles of 256×256 pixels, and generated a binary mask based on the three markers to define the ECM area. We calculated the total per marker expression, normalized by the mask area, for each tile. We discarded tiles that had less than 10% ECM area, and clustered the remaining tiles into Cold Collagen (those with predominantly Collagen expression) and Hot Collagen (those with coexpression of Collagen and Fibronectin and/or FAP). We calculated the proportion of each image that was composed of Cold Collagen, Hot Collagen, or non-ECM tiles.

#### ECM pixel clusters

To capture pixel-level co-expression of ECM markers, we performed pixel clustering on Collagen, Fibronectin, FAP, SMA, and Vimentin using Pixie^28^. We identified a total of 15 distinct ECM pixel clusters, which we used to compute a number of distinct features (**Extended Data Fig. 7h**). We calculated the density and proportion of each cluster in each image as features. For each pixel cluster, we identified groups of contiguous pixels from the same cluster, which were binarized into masks. We then calculated morphological features of these pixel cluster masks to capture their shape. We calculated the per-cluster average of these morphological features for each cluster in each image, which we included as additional features. We then defined pixel neighborhoods using k-means clustering, finding five distinct ECM neighborhoods. We calculated the density and proportion of each neighborhood in each image as features.

#### Aggregating computed features

After computing each of the above image features, we aggregated them into a single data structure for downstream analysis. Each feature was given an informative name, as well as metadata relating to which image compartment it was calculated in, the cell types and/or markers used to calculate the feature, and the broad feature category it belonged to (**Supplementary Table 4**). For features that were in compartments, we assessed the correlation between the compartment-specific value and the image-wide value. Compartment features that had a greater than 0.8 correlation with the corresponding image-wide value were excluded. We then Z-scored each feature to enable easy comparison across feature types. Finally, for samples with multiple FOVs per timpeoint, we computed the average across all the distinct FOVs to use for downstream analysis.

#### Assessing pipeline robustness

To determine the impact of the key design decisions in our pipeline, we systematically varied different thresholds and cutoffs to understand their impact on the output. In general, we found that small changes to any of the parameters resulted in negligible changes, whereas larger (log_2_ ratio of at least 1) produced a noticeable impact on the results. We first looked at the minimum cells per image used for calculating per-cell statistics. In line with the different abundance of different cell types, we observed varying baseline levels of exclusion across distinct cell types (**Extended Data Fig. 7i**). Altering this threshold (which is set at a five cell minimum) affected the number of FOVs excluded, with changes roughly proportional to the change in threshold. We next looked at the functional marker positivity thresholds (**Extended Data Fig. 6l**). We observed relatively minor shifts in total positive cells for small relative shifts in the threshold, with the most significant changes for very low values of the thresholding, which resulted in many cells being called positive (**Extended Data Fig. 7j**). We then looked at the compartment masks, and analyzed how altering the radius of the border region impacted border size. We found a consistent relationship between border radius and total border area (**Extended Data Fig. 7k**), with the most significant deviations only occurring for large (log_2_ ratio of at least 1) changes in the radius.

### Sequencing data

#### DNA and RNA data generation

We used the previously generated DNA and RNA data from all stage 1 patients, which was previously described in Voorwerk et al^24,25^. In addition to the original 60 patients described in that study, we analyzed sequencing data from an additional 33 patients.

#### DNA sequencing processing

To ensure the computational reproducibility, harmonizing and processing the genomic data from these large cohorts at scale, we have deployed Isabl platform locally and developed containerized and version control applications. We integrated tools for identification of Single Nucleotide Variants (SNVs), Copy Number Alterations (CNAs), HLA typing, Neoantigen prediction, IC-subtype prediction and RNA quantification that are described below. For seamless integration of these tools with the Isabel platform, Docker containers for all relevant tools and algorithms are openly available on Docker Hubs (https://hub.docker.com/u/cancersysbio, https://hub.docker.com/u/asntech).

Starting from FASTQ files, we aligned paired whole-exome sequences (n=78 tumor-normal pairs) to the human reference sequence (GRCh38, GATK version downloaded from AWS iGenome, https://ewels.github.io/AWS-iGenomes/) using BWA-MEM (v0.7.17)^74^ as implemented in the TCGA-ICGC-PanCancer PCAP-core docker (https://github.com/cancerit/PCAP-core, v5.6.1). We assessed data quality using FastQC (http://www.bioinformatics.babraham.ac.uk/projects/fastqc/, v0.11.9) and Qualimap (v2.2.1)^75^. Median tumor coverage assessed by Qualimap was 190.0 and 81.1 for tumor and normal samples, respectively.

We detected SNVs using a consensus approach leveraging two independent callers: Mutect2 (v4.1.7.0)^76^ and Strelka2 (v2.9.10)^77^. We ran Mutect2 on tumor/normal pairs as part of the nf-core/sarek Nextflow (v20.12.0) pipeline with the following parameters: -r 2.7.1 –step variantcalling –skip_qc all –tools Mutect2 -profile singularity. We ran Strelka2 on tumor/normal pairs with –exome and –callRegions options with the target bed file. We also leveraged indels called by MANTA (v1.6.0) using the parameters –indelCandidates. A consensus variant call set was obtained by combining Mutect2 and Strelka2 variants using ‘VariantFilter’ (https://github.com/rschenck/VariantFilter). Mutational signatures of SNVs were calculated by deconstructSig R package (v1.8.0)^78^.

We estimated tumor cell fraction and mean ploidy and found allele-specific copy number aberrations using FACETS SUITE (https://github.com/mskcc/facets-suite, v2.0.8) with an allele-specific CNA caller FACETS (v0.6.1)^79^. First, we generated snp pileup files using snp-pileup-wrapper.R using dbSNP (v138) and default parameters. Next, we ran run-facets-wrapper.R using the following parameters: –purity-cval 600 –cval 300 –normal-depth 40. Loss of heterozygosity (LOH) of HLA genes was defined if the sample has a zero minor copy number in HLA regions (chr6: 29.9 - 31.4Mbps). We used iC10 (R package v1.5) with both RNA and DNA-sequencing data (iC10 DNA+RNA) to predict the integrative clustering (IC) subtypes of breast cancer. We also used EniClust (Houlahan, Mangiante, Sotomayor-Vivas, Adimoelja et al., in revision), which reliably infers IC subtypes from DNA-sequencing data of lesions from different stages of progression and diverse organs (primary and metastatic). To ensure sufficient representation of all subtypes when calling the IC10 subtypes, TONIC was integrated with 1102 breast cancer samples from TCGA as a reference panel. RNA from the two datasets was normalized together to remove batch effects using the variance stabilizing transformation (VST) implemented in DESeq2 (v1.26.0).

HLA typing was performed using Optitype (v1.3.5)^80^. We extracted reads mapping to chromosome 6 and remapped them to HLA reference fasta provided by Optitype using BWA-MEM. We then back-extracted mapped reads using fastq function in samtools (v1.15.1)^81^ and used them as input to OptiTypePipeline.py using default parameters. Leveraging somatic SNVs and HLA genotypes, we perform antigen prediction on each sample using antigen.garnish R package run via docker andrewrech/antigen.garnish:2.3.1^82^. We filtered the initial list of putative neoantigens based on default thresholds of foreignness score and agretopicity score, as well as less than 1000nM for binding strength. We called clonal neoantigens using the ccf-annotate-maf function from facetsSuite R package (version 1.0). We categorized neoantigens as clonal if the corresponding SNV was annotated as clonal, and subclonal if otherwise.

#### RNA sequencing data

We aligned raw reads to reference GRCh38 using STAR (v2.7.9a) and read counts were estimated using RSEM (v1.3.3) and GTF file from GENCODE v39. We calculated transcripts per million (TPM) values using expected read counts and effective lengths of protein-coding genes from RSEM. The cytolytic activity (CA) score was calculated by the geometric mean of TPM values of GZMA and PRF1^46^. T cell-inflamed gene expression profile (GEP) signatures were calculated by the mean of log10TPM values of 18 previously defined genes^83^. We followed the TME classification procedure described in the previous study^84^ to define TME subtypes. We scored the 29 signatures defining the TME subtypes, performed median normalization of the resulting scores, and classified them considering the pan-cancer TCGA samples from the original publication as a reference. PAM50 subtypes were computed by genefu R package, integrating TONIC RNA-sequencing with RNA-sequencing of 1102 breast cancer tumors from TCGA to ensure the expected representation of all breast cancer subtypes. The two datasets were normalized using the variance stabilizing transformation (VST) implemented in DESeq2 (v1.26.0). We computed additional Kegg and Biocarta signatures for immune cell polarization and cytokine secretion for a total of 134 RNA-based signatures (**Supplementary Table 5**).

#### Feature extraction

Following the generation above the features, we processed them to be used for multivariate modeling. The following features were included: tumor cell fraction, mean ploidy, fraction of copy-number alteration, fraction of LOH, whether a sample had whole genome doubling, IC subtype, arm-level copy number alterations, copy number of individual genes from iC10 package and previous studies^85,86^, number of missense, synonymous, frameshift, clonal, and subclonal mutations, neoantigen burden (clonal and subclonal), SNV signatures, HLA type, HLA-binding mutation ratio as implemented in Van Den Eynden *et al.*^87^, and all 134 RNA signatures.

### Univariate outcome associations

#### Calculating feature associations

To link features with patient outcome, we separately analyzed the MIBI, RNA, and DNA data. We also analyzed each timepoint separately. We used responder/non-responder status (see Study design, above) as the outcome variable, which is a binary classification for whether a given patient did, or did not, respond to IC from the trial. For each feature, we calculated an independent t-test to determine if there was a difference in means between the two populations. We recorded the p-value for each comparison, as well as the difference in medians between the two populations.

#### Ranking features

To determine the relative importance of each feature, we ranked each p-value from smallest to largest. We then ranked the shifts in the median from largest to smallest. We averaged these ranks together for each feature, to arrive at a composite metric, the importance score. We generated a separate set of importance scores for the MIBI, RNA, and DNA data. For each modality, we calculated a single set of importance scores across all timepoints to evaluate their relative importance.

#### Assessing importance score robustness

To ensure that the importance score was accurately capturing informative features, we assessed the robustness of the identified features. We randomly shuffled the outcome labels for each patient, and regenerated the univariate outcome association metrics. We then compared the top 100 features in the real data with the top 100 features in the shuffled dataset, using the importance score to identify the top 100 separately in each dataset. Looking at the distribution of p-values in the real data compared to one of the shuffled replicates, we see that the distribution of the real data is shifted to the right with almost no overlap, indicating that the real data has much more significant p-values (**Extended Data Fig. 9a**). We then compared the average p-value in the real data with the average p-value in each of the 100 shuffled iterations, finding that the real data is completely non-overlapping with the observed distribution of shuffled averages (**Extended Data Fig. 9b**).

We performed the same analysis looking at the difference in medians between the two populations, as this is the second metric that is used to define the importance score. Looking at the distribution of medians from the real data and an individual permutation, we see a rightward shift in the medians, though not as extreme as was observed for the p-values (**Extended Data Fig. 9c**). Looking at the averages across all the replicates, we see that the observed average is almost completely non-overlapping with the randomized averages (**Extended Data Fig. 9d**), indicating that we are identifying features with much larger effect sizes and much smaller p-values than expected.

To determine if there were certain features which were more likely than others to come up as false positives, we looked at the specific features selected as part of the top 100 in each of the above replicates (**Extended Data Fig. 9e**). Overall, we observed relatively little overlap between the same features across distinct replicates, suggesting that there were not underlying factors influencing the features which we identified. We also looked at the correlation of the features selected as part of the actual top 100 compared to the rest of the dataset (**Extended Data Fig. 9f**). These features were more correlated with one another than with the non-selected features, which makes sense given the shared patterns in the types of response-associated features we identified.

#### Ranking evolution features

The features above were all determined by looking exclusively at a single, static timepoint. We took advantage of the paired timepoints in our cohort (**Fig. 1a**) to calculate the change in each feature from primary to baseline, baseline to pre-nivo, baseline to on-nivo, and pre-nivo to on-nivo. We then repeated the same analysis as above, ranking these features based on their association with patient outcome. Looking at the top features across both static and evolutionary features, we observed substantial overlap, with many of the same features appearing in both analyses (**Extended Data Fig. 9g**). We therefore focused the analyses in the paper on the static features, which do not require paired samples to calculate.

### Multivariate modeling

To predict patient response to treatment, we used a Lasso^42^ model, which is well-suited for handling high-dimensional datasets with many features and relatively few data points. The Lasso model’s ability to select the most relevant features enhances its predictive performance in such scenarios. We also experimented with other models, including XGBoost, but found that the Lasso model demonstrated overall better effectiveness given our dataset’s structure.

Specifically, we utilized stratified cross-validation (CV) to select the level of sparsity in the Lasso model (**Extended Data Fig. 12a)**. As compared to the regular CV procedure, stratified CV maintains a balanced representation of patient responses in each fold. This is achieved by leveraging the available patient response information as a covariate during the fold-splitting process. By stratifying the folds based on the response covariate, we ensure that each fold contains a representative sample of both responders and non-responders, mitigating the potential bias that could arise from an imbalanced distribution. We used a three-fold CV in our experiments. Moreover, in order to draw more robust conclusions, we conducted the same set of experiments 10 times with different random seeds. The Area Under the Receiver Operating Characteristic Curve (AUROC) was used as the primary metric for evaluating model performance.

To gain insights into the model’s decision-making process, we examined the weights assigned by the Lasso model to each feature. The Lasso model’s sparse solution results in many feature weights being set to zero, effectively performing feature selection. By analyzing the non-zero feature weights, ranked by their absolute magnitude, we can identify the most important features contributing to the model’s predictions. The magnitude of the non-zero weights provides a measure of the relative importance of each selected feature, with larger absolute values indicating a stronger influence on the model’s predictions of patient response.

To identify the features most important for the model’s prediction, we defined a set of top features across the 10 replicates. In order for a feature to be considered a top feature, it needed to be selected by the model in at least three separate replicates, and it needed to have a weight of at least 30% as large as the largest feature. Using this criteria, we sought to understand how consistently top features were identified across distinct replicates. We found that more than half of the top features were identified in at least 80% of trials, indicating good consistency across the distinct splits (**Extended Data Fig. 12b**).

One of the reasons to use a Lasso model is that the output is sparse, meaning only a small subset of the features are included. As a result, the selected features will tend not to be correlated with one another. As expected, the top features selected by the model were less correlated with another than the top 100 features selected by the univariate analysis (**Extended Data Fig. 12c**). We next compared the univariate scores of the top features selected by the model. For the on-nivo MIBI model, the top features had very high importance scores (**Extended Data Fig. 12d**). The on-nivo RNA model had one top feature with high importance score (the cytolytic activity score), and one feature with a relatively low importance score (anti-tumor cytokines), indicating that the model prioritized different information from the univariate analysis. In agreement with the relatively small degree of sharing across timepoints for features identified in the univariate analysis (**Fig. 4b**), top features from the models likewise were not often shared across timepoints (**Extended Data Fig. 12e-f**).

Our analysis showed the on-nivo timepoint as the most predictive. To further evaluate the predictive capability and generalizability of this model, we divided the dataset using stratified splitting into training and test sets, with a 70% (48 patients) and 30% (21 patients) ratio. We standardized the training set and applied the derived mean and standard deviation to the test set to prevent data leakage. The optimal level of sparsity in the Lasso model was determined through stratified CV within the training set. The model’s performance was then evaluated on the unseen test set, with results reported in terms of both the AUROC and Area Under the Precision-Recall Curve (AUPRC) (**Extended Data Fig. 12g-h**). We observed similar performance with this setup (AUROC=0.875) compared with the cross-validation framework we used for the preceding analyses, indicating that the performance metrics we used were unlikely to be inflated by data leakage.

## Data availability

Sequencing data and source data supporting the findings of this study will be made available upon reasonable request for academic use, within limitations of the provided informed consent. RNA and DNA-sequencing data on tumor biopsies of TNBC patients treated in the TONIC-1-trial stage 1 are deposited at the European Genome-phenome Archive (EGA) under accession number EGAS0001003535 RNAseq data of TNBC patients treated in TONIC-1-trial stage 2 reported in the paper are not deposited in a public repository pending ongoing work but can be made available from the corresponding authors upon reasonable request. All human data requests will be reviewed by the Institutional Review Board (IRB) of the NKI and applying researchers have to sign a data transfer agreement after IRB approval before the data can be released (contact: co-last author M.K.).

The MIBI image data is available for browsing at: https://idr.openmicroscopy.org/webclient/?show=project-3052

The MIBI image data as well as corresponding image masks can be downloaded at: https://www.ebi.ac.uk/biostudies/bioimages/studies/S-BIAD1288

A subset of the processed image data is available in an interactive viewer, which can be accessed at: https://angelolab.github.io/tonic-vitessce/

All analysis files can be accessed here: https://zenodo.org/records/14112853

## Code availability

The code to generate the figures in this paper is available at: https://github.com/angelolab/publications/tree/main/2024-Greenwald_Nederlof_etal_TONIC The low-level processing code is available at: https://github.com/angelolab/toffy

The segmentation and cell assignment pipelines are available at: https://github.com/angelolab/ark

SpaceCat is available at: https://github.com/angelolab/SpaceCat

## Author contributions

N.F.G., I.N., C.C., M.A., and M.K. conceived the project.

I.N. collected the longitudinal samples and clinical data.

M.d.G. collaborated on clinical data collection

L.V. and V.G. coordinated TONIC trial procedures.

M.d.M. processed FFPE for IHC, isolated DNA and RNA from tissue biopsies, constructed the tissue microarrays.

M.J.vdV and H.M.H. annotated FFPE tissue for TMA construction and provided a pathology review of each tissue slide.

M.J.vd.V funded longitudinal sampling and the logistical part of the project.

T.N.S provided guidance.

N.F.G., C.C.F., Z.K., Y.B., E.M., H.P., T.R., A.D., F.H., and M.B. generated the MIBI data

N.F.G., C.S., D.D., A.Ko., I.N., S.R.V., E.M., C.C.L., J.R., A.Ka., and R.T. processed and analyzed the MIBI data

S.P., K.H., A.Kh., N.F.G., D.D., C.C.L, C.Y.Y., L.M., C.S-V., and Z.M. processed and analyzed the RNA and DNA sequencing data

N.F.G., I.N., C.S., D.D., S.P., A.Ko., and S.R.V. generated the figures

N.F.G., I.N., C.C., M.K. and M.A. wrote the manuscript.

C.C., M.K. and M.A supervised the project.

All authors provided feedback on the manuscript.

## Acknowledgements and funding

We thank the patients and their families for participating in the TONIC trials. TONIC trial costs were supported by Bristol Myers Squibb. The funders had no role in study design, data collection or analysis, decision to publish or preparation of the manuscript. We thank all supporting clinical trial staff, in particular nurse specialists and the Departments of Medical Oncology, Surgery, Radiology and Pathology of the participating centers. We thank the NKI Core Facility of Molecular Pathology & Biobanking for their support in processing of samples. Support for title page creation and format was provided by AuthorArranger, a tool developed at the National Cancer Institute. Figures and graphics were created with Biorender.com. We thank T.M. for constant support.

N.F.G. was supported by NCI CA246880, NCI CA264307, and the Stanford Graduate Fellowship. L.M. was supported by the Stanford School of Medicine Dean’s Fellowship. Collection and processing of samples was made possible by KWF grant 2016-10510 from the Dutch Cancer Foundation. Research in the laboratory of M.K. is funded by the Netherlands Organization for Scientific Research (VIDI) and Victoria’s Secret Global Fund for Women’s Cancers Rising Innovator Research Grant, in Partnership with Pelotonia & AACR.

