## Extended Data for "Temporal and spatial composition of the tumor microenvironment predicts response to immune checkpoint inhibition"

#### Extended Data Figure 1. ROI selection and sample summary

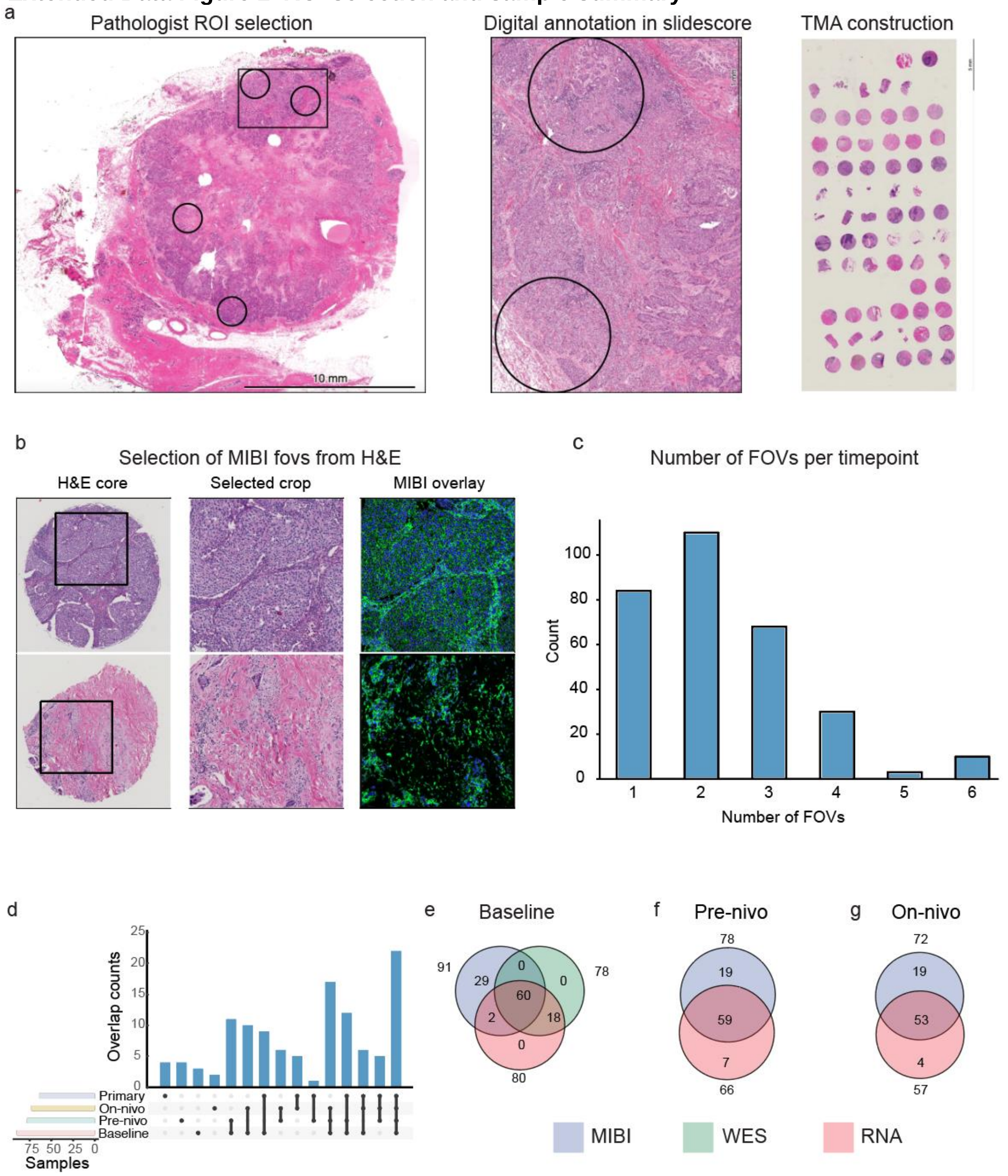


**a**) Workflow for selecting cores to be included in the study. Hematoxylin and eosin (H&E) slides were digitized in slidescore.com, annotated by a dedicated breast cancer pathologist, and demarcated with areas of interest. Each identified region was punched with a 1.5mm core and placed onto a tissue microarray (TMA). **b**) Illustrative examples of the H&E image of the entire core, the cropped inset of the portion of the core selected for MIBI analysis, and the corresponding MIBI image generated from the selected region. **c**) Histogram showing the number of fields-of-view (x-axis) acquired on MIBI per tumor sample (y-axis). **d**) Upset plot showing the overlap of distinct timepoints of MIBI data across the patients in the cohort. **e**) Venn diagram showing the overlap between DNA, RNA, and MIBI data for baseline samples. **f**) Venn diagram showing the overlap between RNA and MIBI data for pre-nivo samples. **g**) Same as f) for on-nivo

#### Extended Data Figure 2. Antibody panel validation

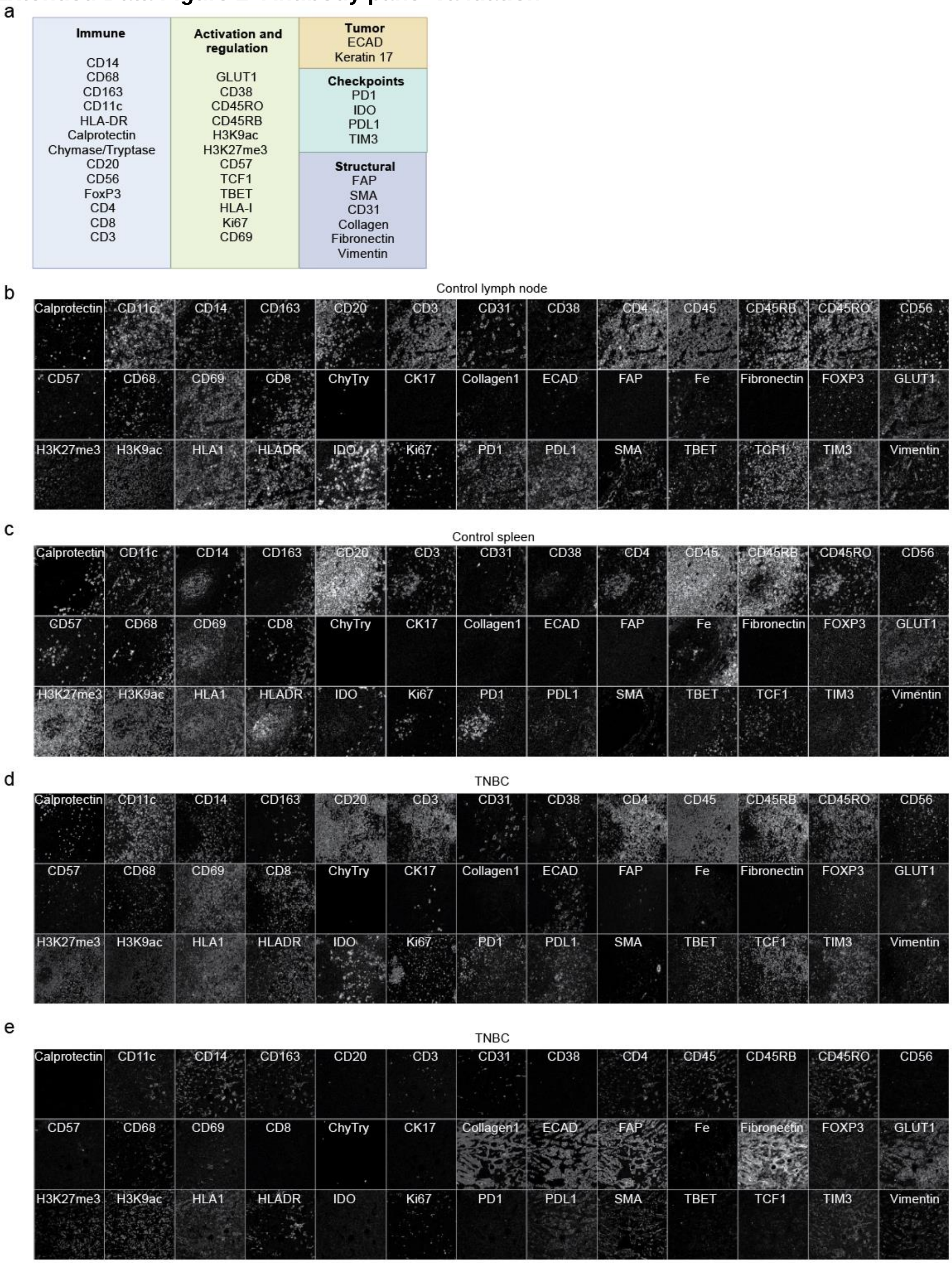


**a**) MIBI panel, including immune markers, tumor markers, checkpoint markers, structural markers and markers related to activation and regulation. **b**) Single-plex images of each marker in the panel in a control lymph node sample. **c**) Same as b) for a control spleen sample. **d**) Same as b) for first representative TNBC sample. **e**) Same as b) for second representative TNBC sample.

#### Extended Data Figure 3. Rosetta image compensation

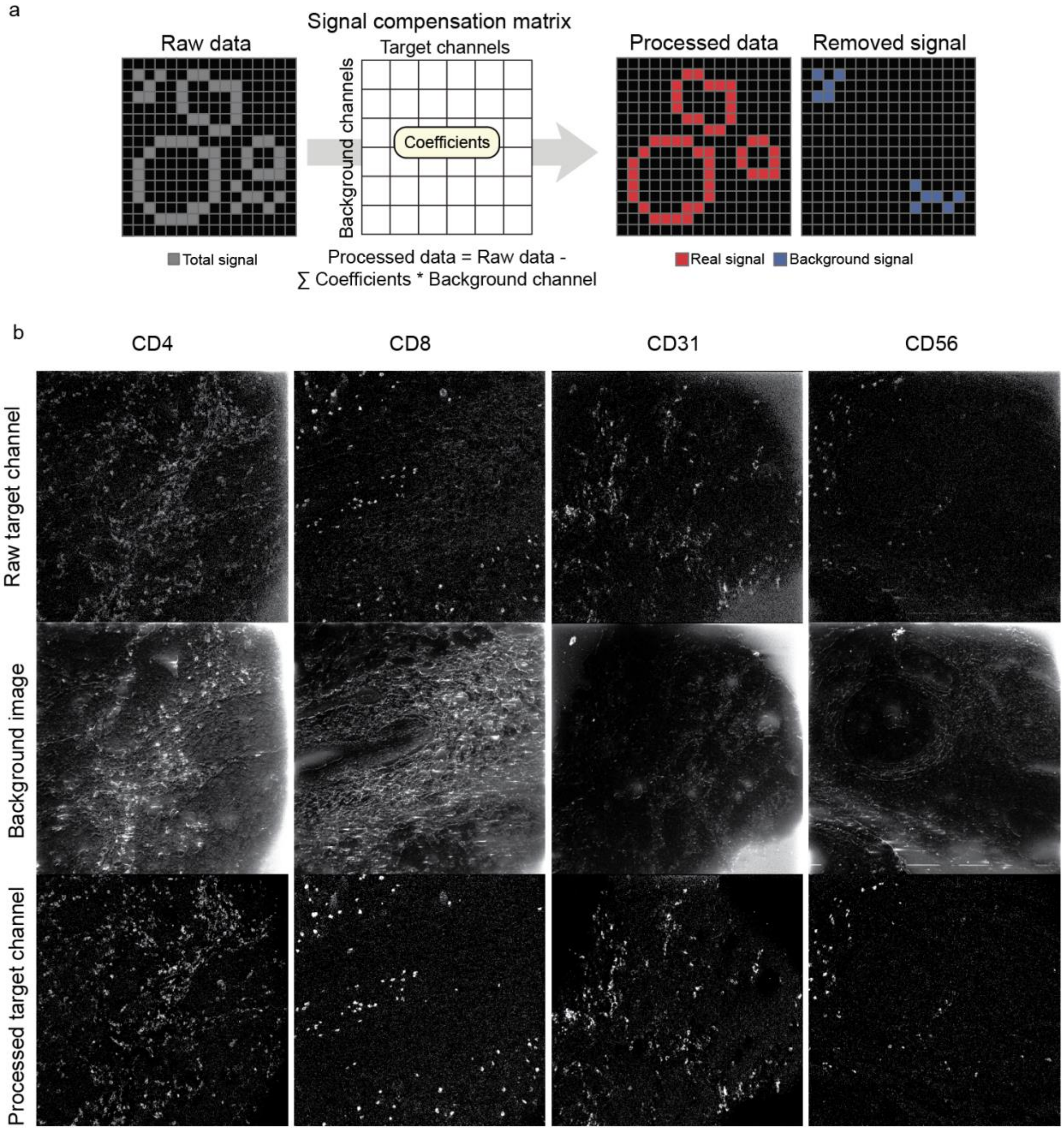


**a**) Cartoon illustrating the Rosetta image compensation process, where background signal is subtracted using a compensation matrix to produce cleaned up images. **b**) Representative examples of different channels before compensation (top), after compensation (bottom), along with the corresponding background channel used for compensation (middle) from the same image.

#### Extended Data Figure 4. Image normalization and quality control

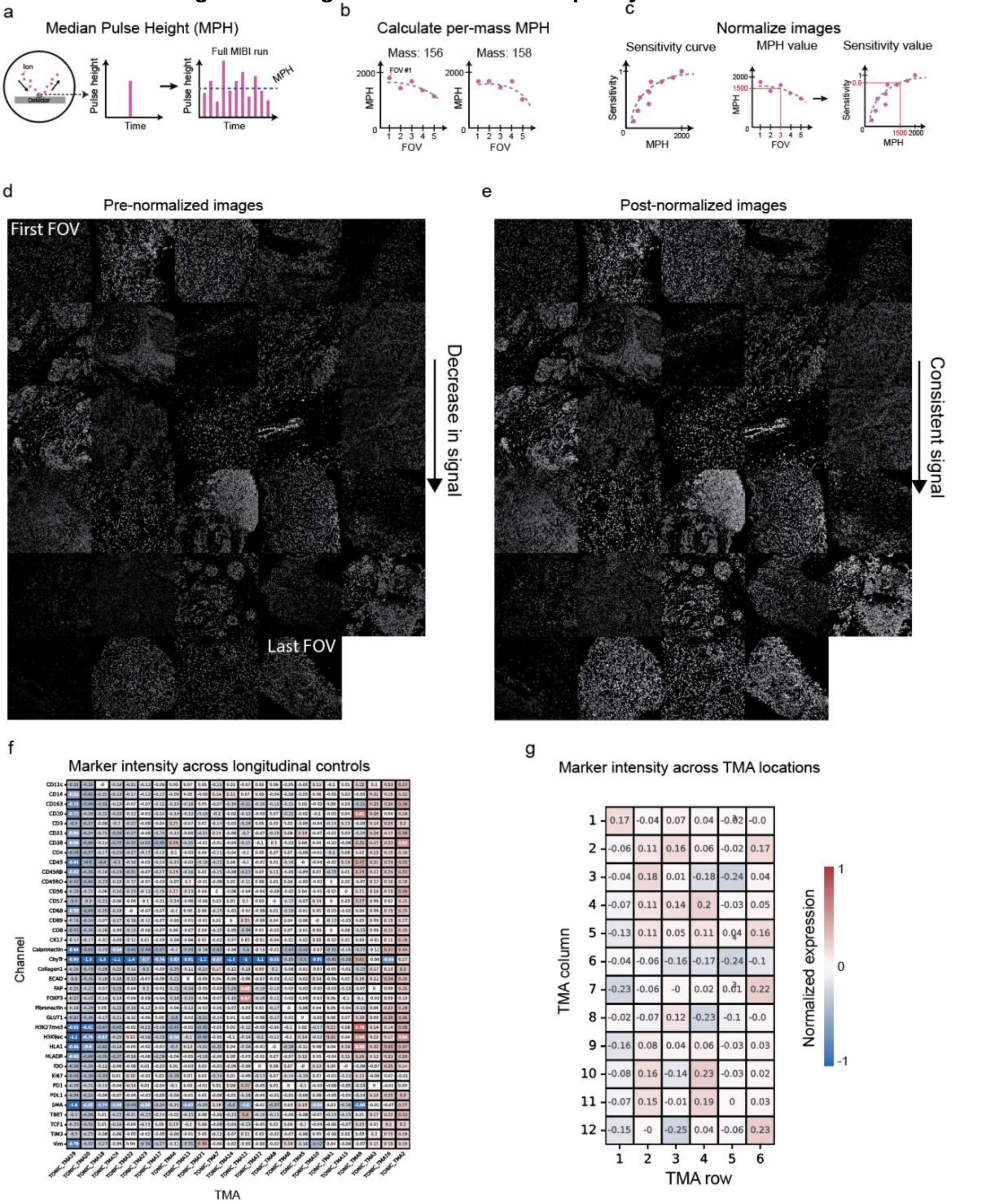


**a**) Cartoon illustrating how median pulse height (MPH) is calculated. **b**) Cartoon showing how a per-mass MPH curve is fit for an imaging run. **c**) Cartoon showing how the sensitivity curve is combined with the per-mass MPH curve to correct for changes in sensitivity over the course of an imaging run. **d**) Representative stitched image showing the uncorrected images from a single channel across an entire run, stitched in acquisition order from the start of the run (top left) to the end of the run (bottom right). **e**) Representative stitched image showing the corrected intensities from the images in d) following MPH normalization. **f**) QC plot showing the normalized expression in the control tissues of all the markers in the panel (y-axis) across the different TMAs in the cohort (x-axis). **g**) QC plot showing the normalized expression per field-of-view, plotted with the same row/column location as in the actual TMA.

#### Extended Data Figure 5. Cell segmentation

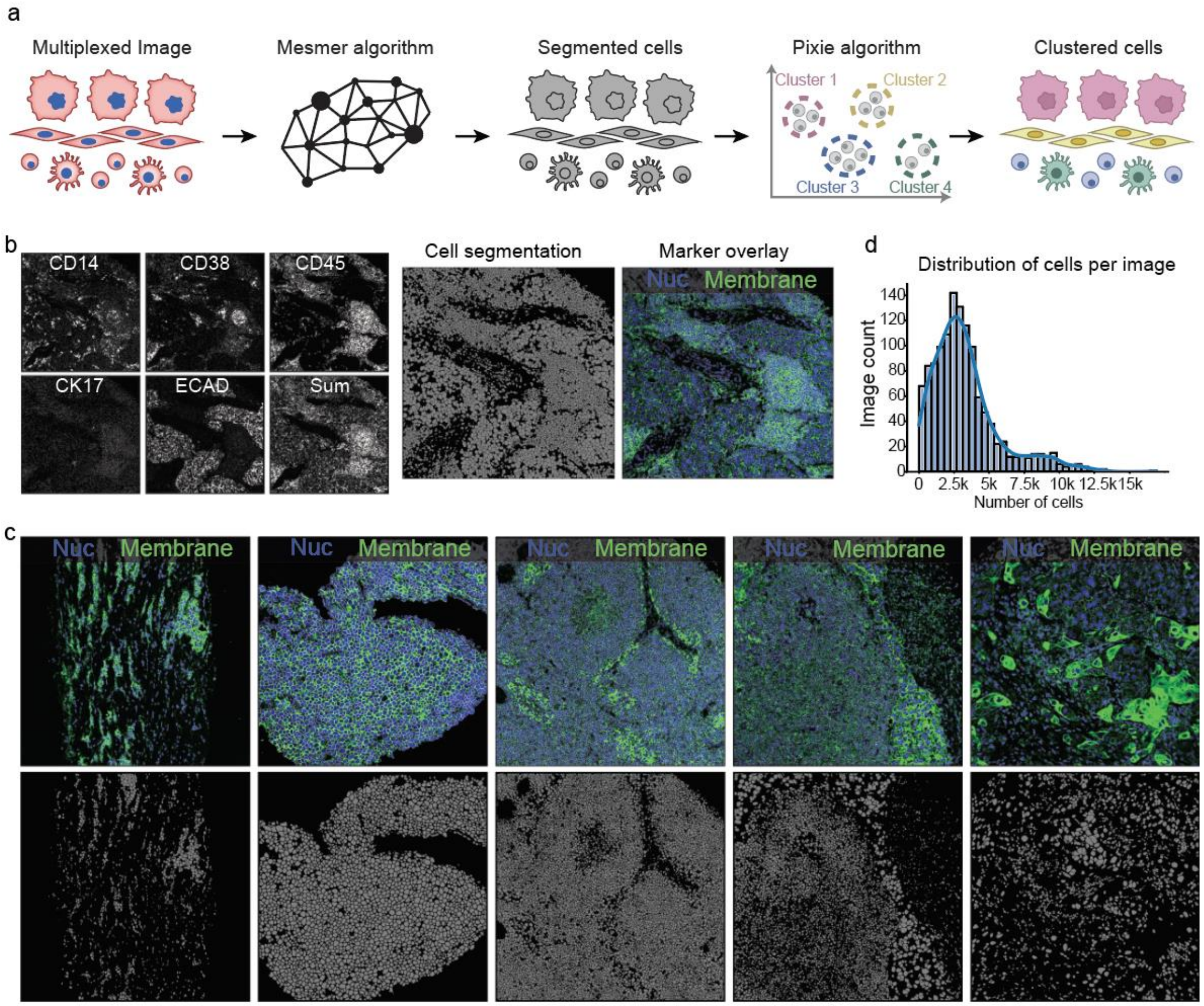


**a**) Cartoon of the segmentation and clustering pipeline. **b**) Representative image showing the different membrane markers that were combined together into a single membrane channel for segmentation (left), the resulting segmentation mask generated by Mesmer (middle), along with an overlay of segmentation mask and channels used for segmentation (right). **c**) Additional representative images of showing the segmentation channels overlaid with the segmentation outlines (top), along with the segmentation mask on its own (bottom). **d**) Histogram showing the number of cells per image across all images in the cohort.

### Extended Data Figure 6. Cell clustering and abundance

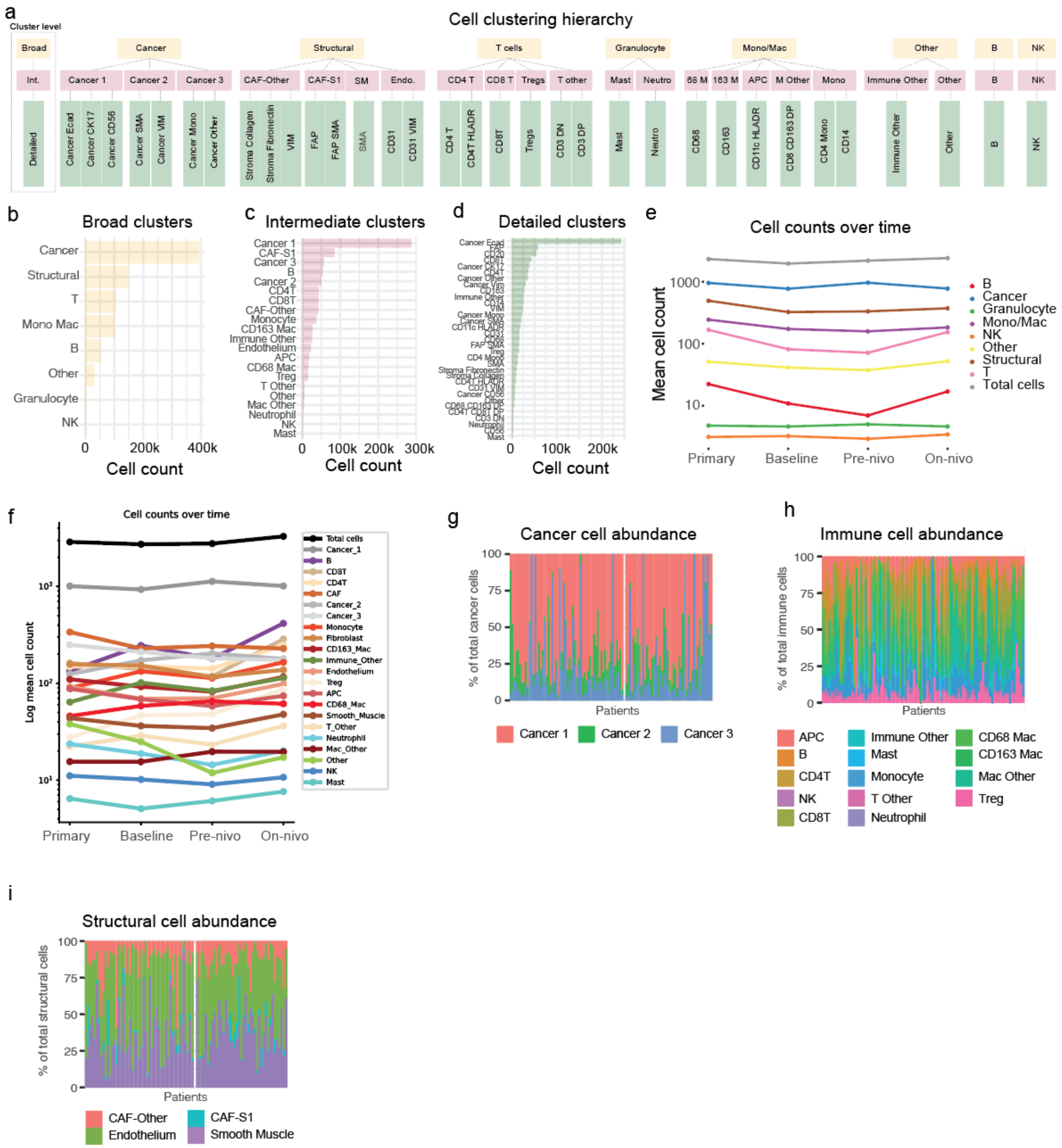


**a**) Diagram illustrating the three levels of cell clustering (broad, intermediate, detailed), and the relationship between clusters in each level. **b**) Number of cells of each cell type in the broad cluster. **c**) Number of cells of each cell type in the intermediate clusters. **d**) Number of cells of each cell type in the detailed clusterings. b, c, d are all based on all primary and metastatic tumors in the sample set. **e**) Mean count of cell types (broad clusters) for each timepoint. **f**) Mean count of cell types (intermediate clusters) for each timepoint. **g**) Average proportion of each intermediate cancer subpopulation in metastatic samples. **h**) Average proportion of each intermediate immune subpopulation in metastatic samples. **i**) Average proportion of each intermediate structural subpopulation in metastatic samples

### Extended Data Figure 7. Feature extraction pipeline cartoons

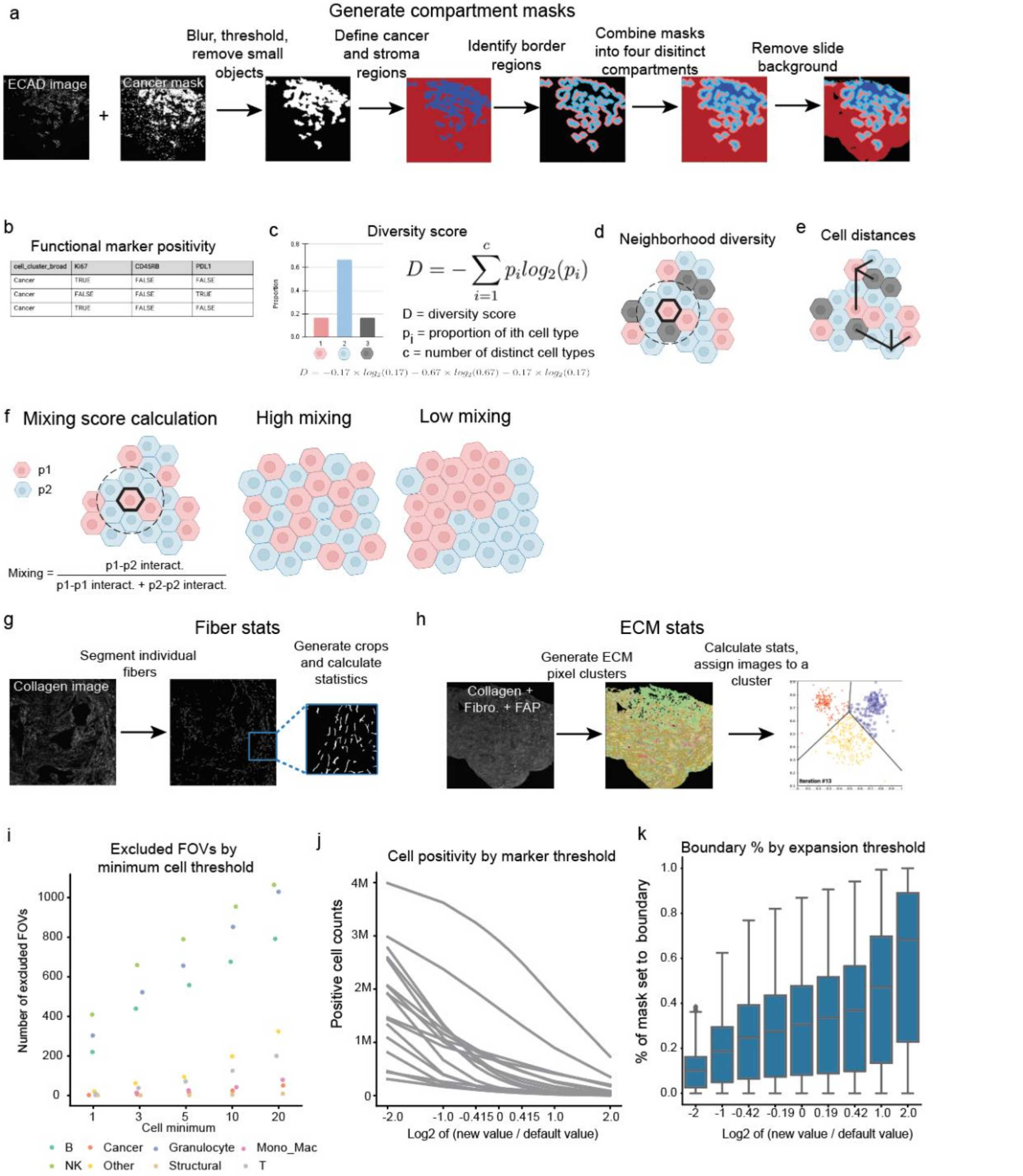


**a**) Cartoon illustrating how compartment masks are defined. **b**) Table illustrating how functional marker thresholds are used to binarize cells into positive or negative for each marker; these assignments are then used to generate positivity proportion statistics per image. **c**) Schematic showcasing how diversity scores are calculated. **d**) Cartoon showing how the radius around a cell is used for neighborhood diversity. **e**) Cartoon showing how cell distances are used to compute the linear distance feature. **f**) Cartoon showing how the mixing score is calculated, along with examples of high and low mixing. **g**) Schematic showing how the fiber segmentation pipeline is used to generate features. **h**) Schematic showing how the extracellular matrix pipeline is used to generate features. **i**) Number of images removed from analysis as a function of different minimum cell thresholds, stratified by cell type. **j**) The number of functional marker positive cells across different thresholds. **k**) The proportion of the image assigned to the border compartment across different expansion thresholds.

#### Extended Data Figure 8. Quantification of features across tumor compartments

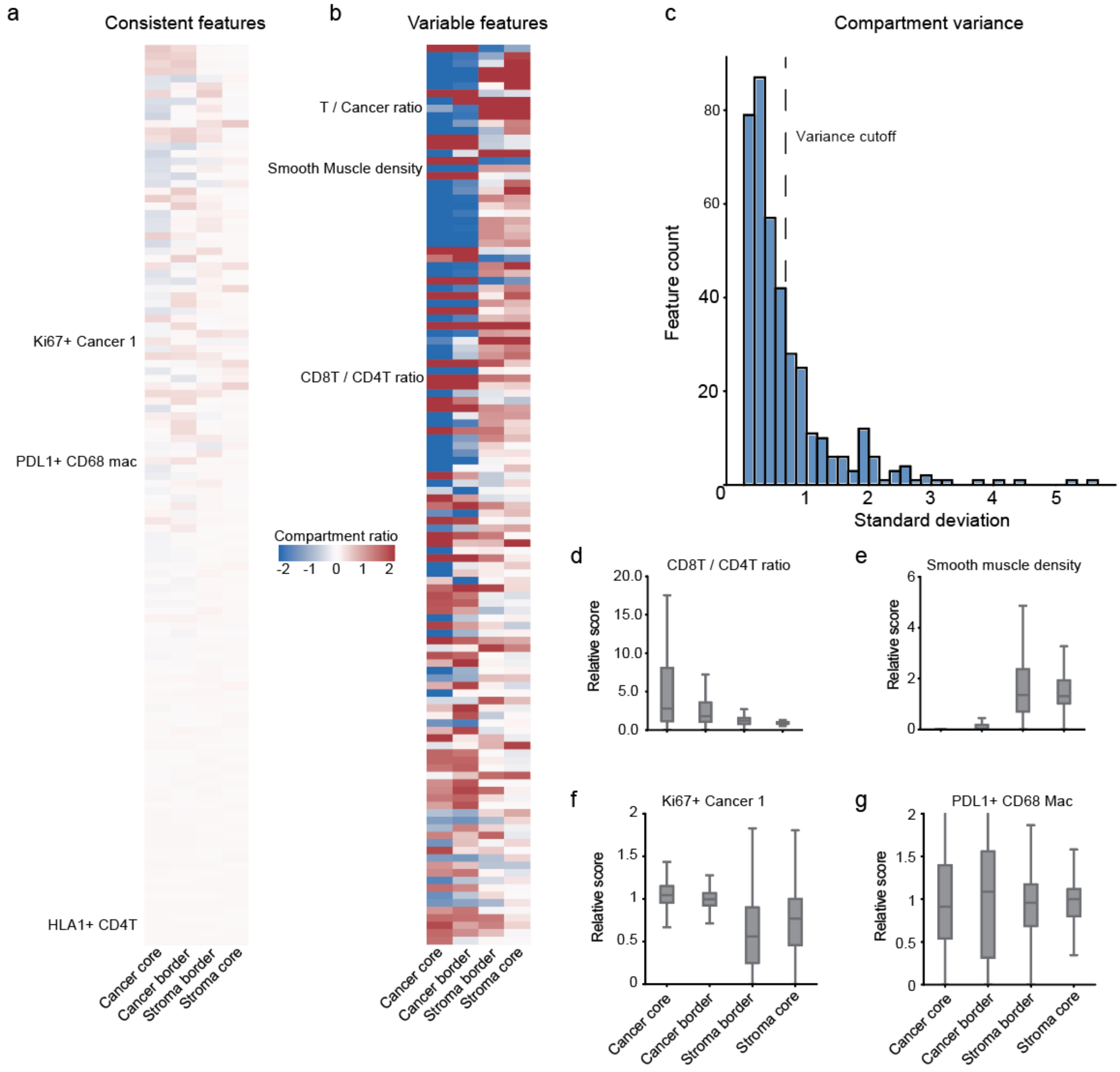


**a**) Heatmap showing features that are consistent across compartments. Each row represents a different feature, and each column is a distinct compartment. Features are normalized to the whole image value. **b**) Same as a), but for features that are enriched in specific compartment(s). **c**) Histogram showing threshold used to identify varying vs. non-varying compartment features. **d**) Distribution of the CD8 T / CD4 T ratio, a feature that changed across compartments. **e**) Distribution of Smooth Muscle density, a feature that changed across compartments. **f**) Distribution of Ki67 positivity in Cancer 1, a feature that did not change across compartments. **g**) Distribution of PD-L1 positivity in CD68 macrophages, a feature that did not change across compartments.

#### Extended Data Figure 9. Univariate outcomes pipeline robustness

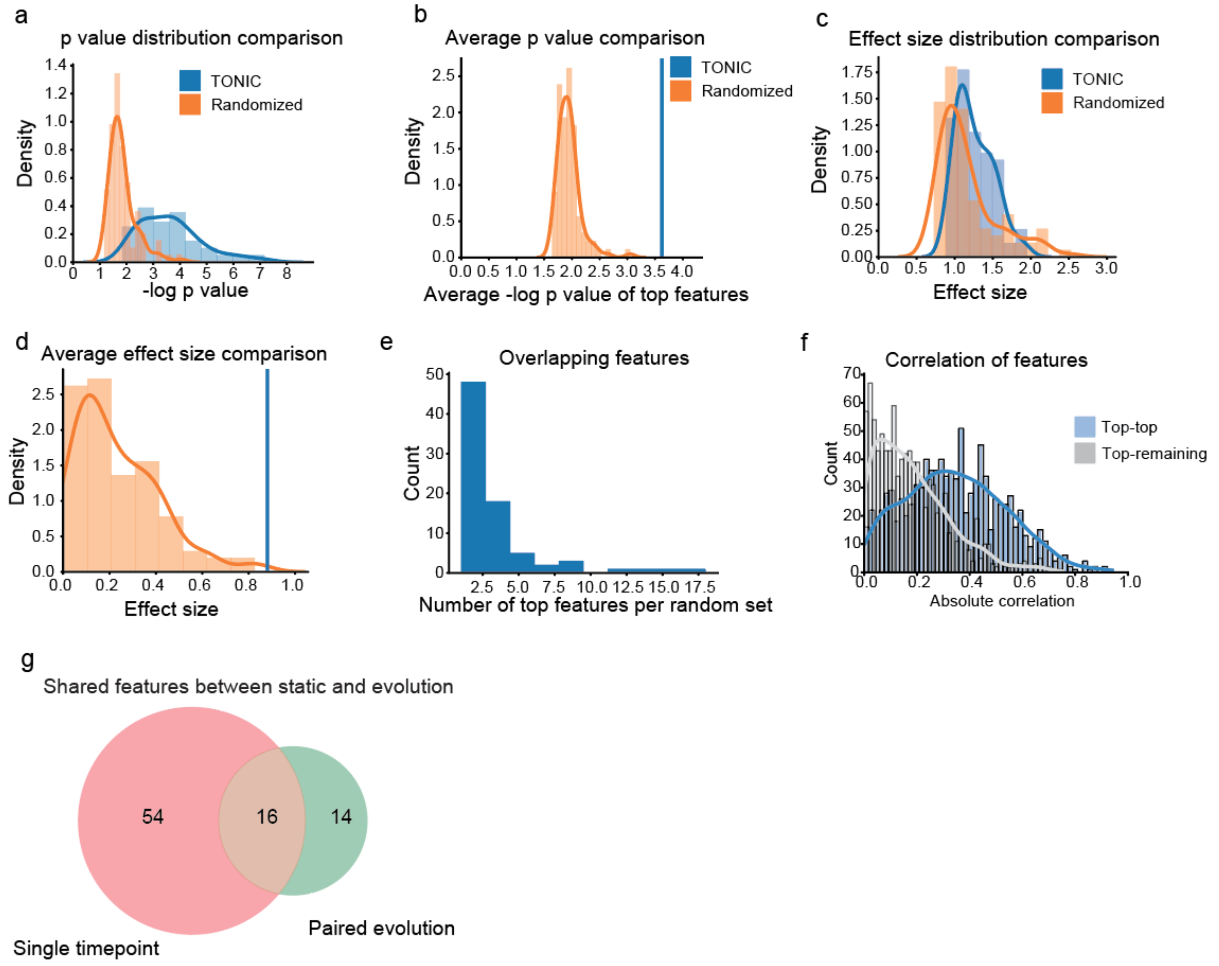


**a**) Distribution of p-values from the top features 100, as well as distribution of p-values from the top 100 features recalculated after the labels have been randomized. **b**) Average p-value in the top features, as well as distribution of average p-values across 100 different randomizations. **c**) Distribution of effect sizes from the top 100 features, as well as distribution of effect sizes from top 100 features recalculated after the labels have been randomized. **d**) Average effect size in the top features, as well as distribution of average effect sizes across 100 different randomizations. **e**) Number of times the same feature come up in different randomizations. **f**) Correlation between top features, compared with correlation between top features and other features. **g**) Overlap in features identified when examining each timepoint independently, compared to looking at the change in each feature across timepoints.

### Extended Data Figure 10. Outcomes associations

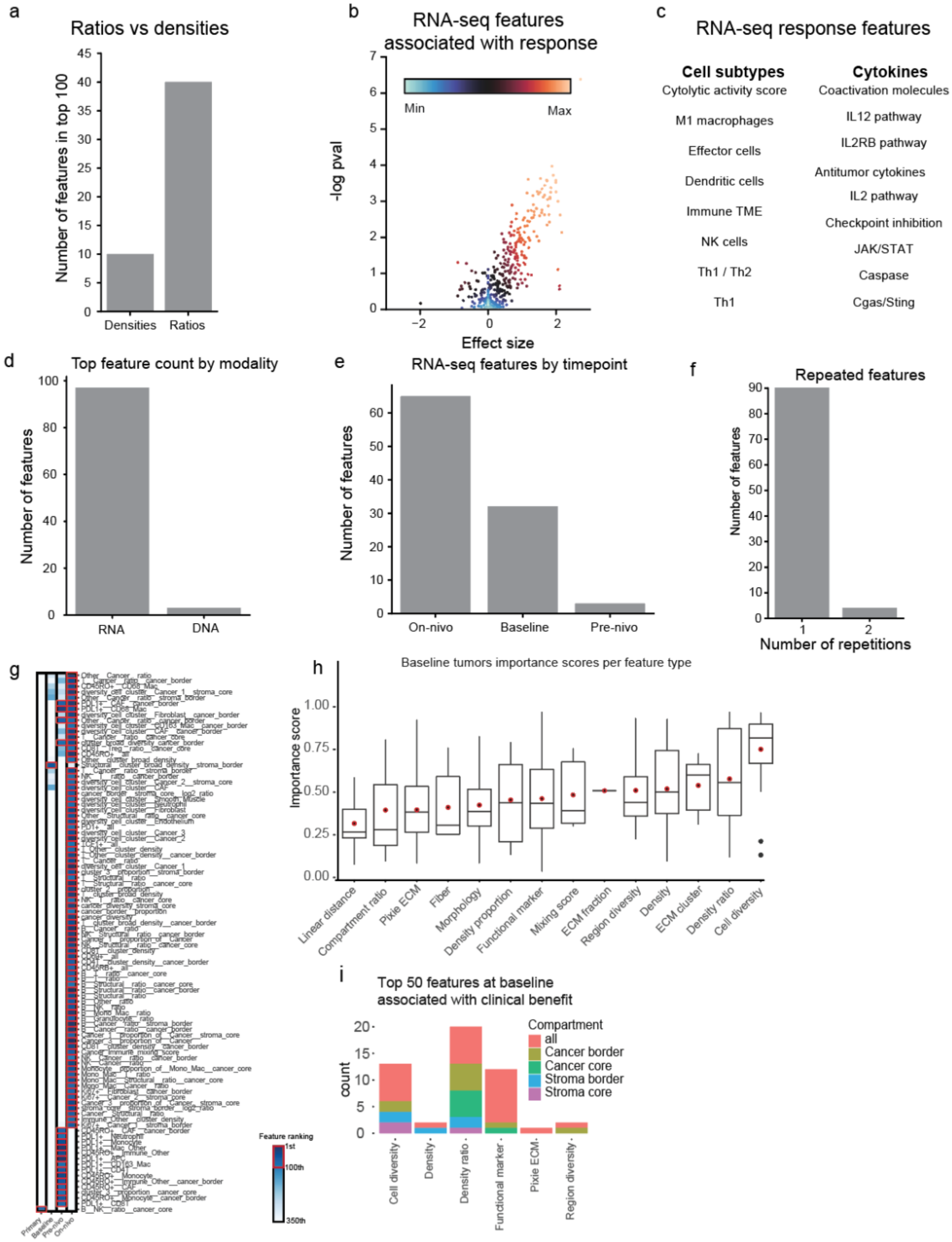


**a**) Enumeration of the number of top MIBI features which are ratios compared to the number which are densities. **b**) Volcano plot of RNA-seq features associated with response. **c**) Top RNA features associated with response, organized by biological category. **d**) Comparison of number of DNA and RNA features associated with response. **e**) Number of RNA-seq features associated with response by timepoint. **f**) The number of times a given MIBI feature was repeated across distinct timepoints among the top 100 features. **g**) Same as Fig. 4b, but with all labels included. **h**) Types of features associated with response from baseline metastatic timepoint. **i**) Top 50 features associated with response from baseline timepoint, colored by compartment and sorted by feature type.

#### Extended Data Figure 11. Timepoint-specific features

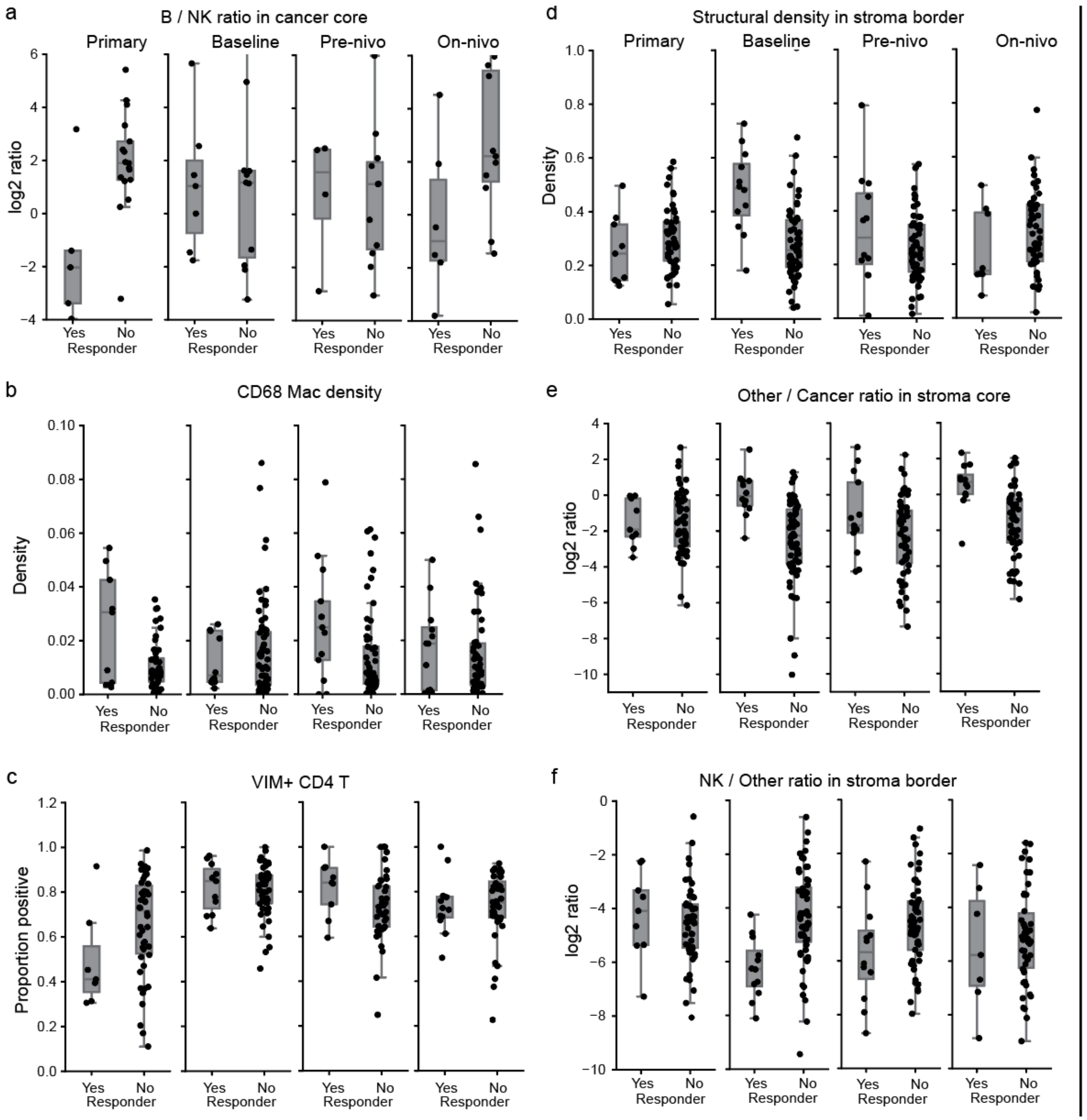


**a**) Box plots showing the distribution of the B cell / NK cell ratio feature in responders and non-responders, across the four distinct timepoints in the cohort. **b**) Same as a), with CD68 macrophage density as the feature. **c**) Same as a), with Vimentin positive CD4 T cells as the feature. **d**) Same as a), with structural cell density in the stroma border as the feature. **e**) Same as a), with the ratio of other to cancer cells as the feature. **f**) Same as a), with the ratio of NK cells to other cells as the feature.

#### Extended Data Figure 12. Multivariate modeling

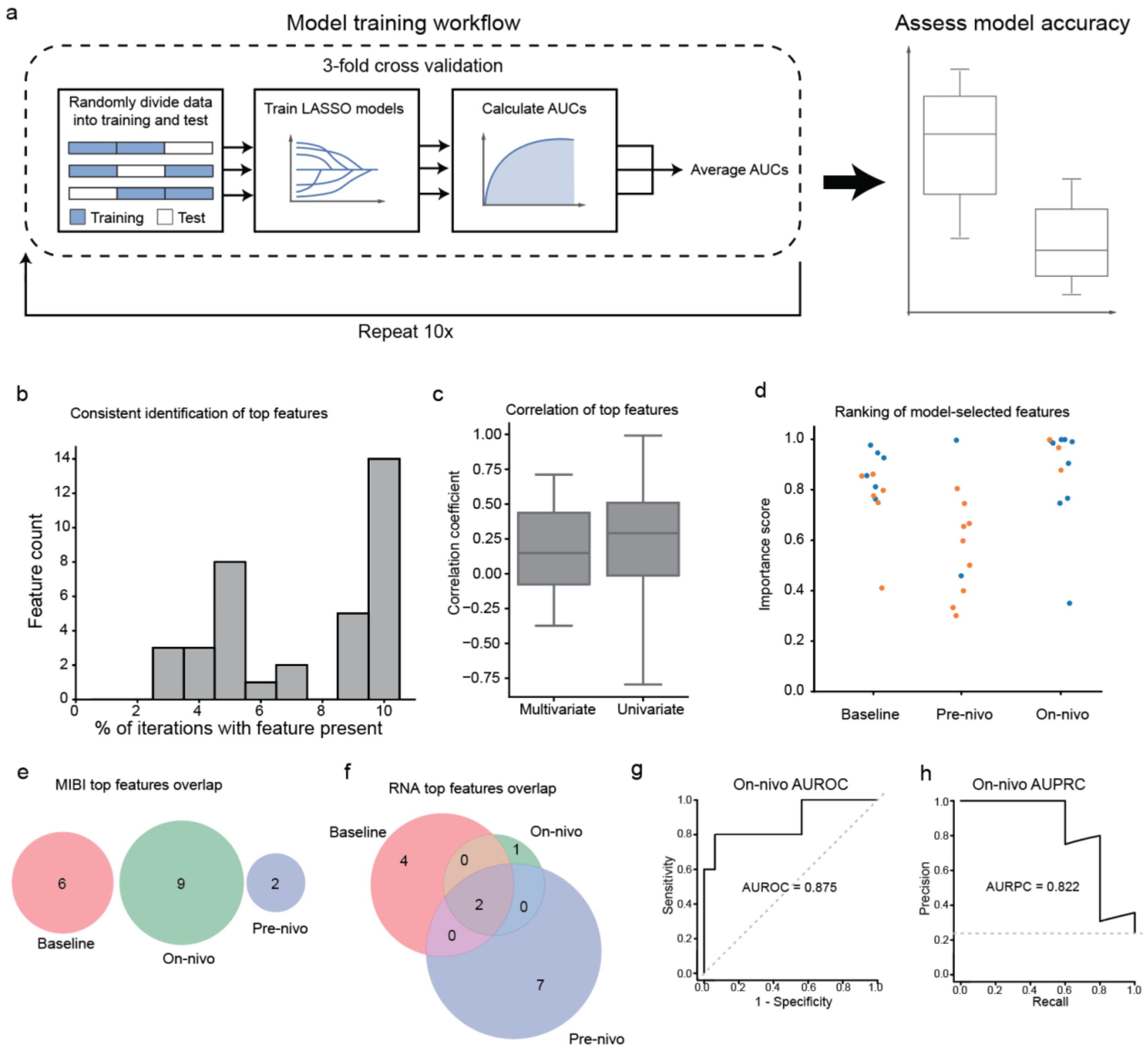


**a**) Diagram illustrating the cross validation approach used for model training. **b**) Histogram showing the number of times the top features were selected across the 10 different iterations of model training. **c**) Comparison between the correlation of the top features identified by the model and the top features from the univariate analysis. **d**) Comparison of the importance score of the top features identified by the model across distinct timepoints, stratified by MIBI vs. RNA. **e**) Overlap across timepoints of top features identified by the MIBI models. **f**) Overlap across timepoints of top features identified by the RNA models. **g**) AUROC evaluated for the on-nivo MIBI data using separate train, val, test split without cross validation. **h**) Same as above, for AUPRC.
